# Wild bird mass mortalities in eastern Canada associated with the Highly Pathogenic Avian Influenza A(H5N1) virus, 2022

**DOI:** 10.1101/2024.01.05.574233

**Authors:** Stephanie Avery-Gomm, Tatsiana Barychka, Matthew English, Robert Ronconi, Sabina I. Wilhelm, Jean-François Rail, Tabatha Cormier, Matthieu Beaumont, Campbell Bowser, Tori V. Burt, Sydney Collins, Steven Duffy, Jolene A. Giacinti, Scott Gilliland, Jean-François Giroux, Carina Gjerdrum, Magella Guillemette, Kathryn E. Hargan, Megan Jones, Andrew Kennedy, Liam Kusalik, Stéphane Lair, Andrew S. Lang, Raphael Lavoie, Christine Lepage, Gretchen McPhail, William A. Montevecchi, Glen J. Parsons, Jennifer F. Provencher, Ishraq Rahman, Gregory J. Robertson, Yannick Seyer, Catherine Soos, Christopher R. E. Ward, Regina Wells, Jordan Wight

## Abstract

In 2022, a severe outbreak of clade 2.3.4.4b Highly Pathogenic Avian Influenza (HPAI) H5N1 virus resulted in unprecedented mortality among wild birds in eastern Canada. Tens of thousands of birds were reported sick or dead, prompting a comprehensive assessment of mortality spanning the breeding season between April 1 and September 30, 2022. Mortality reports were collated from federal, Indigenous, provincial, and municipal agencies, the Canadian Wildlife Health Cooperative, non-governmental organizations, universities, and citizen science platforms. A scenario analysis was conducted to refine mortality estimates, accounting for potential double counts from multiple sources under a range of spatial and temporal overlap. Correcting for double counting, an estimated 40,966 wild birds were reported sick or dead in eastern Canada during the spring and summer of 2022. Seabirds and sea ducks, long-lived species that are slow to recover from perturbations, accounted for 98.7% of reported mortalities. Mortalities were greatest among Northern Gannets *(Morus bassanus*; 26,193), Common Murres (*Uria aalge*; 8,133), and American Common Eiders (*Somateria mollissima dresseri;* 1,945), however, these figures underestimate total mortality as they exclude unreported deaths on land and at sea. In addition to presenting mortality estimates, we compare mortalities with known population sizes and trends and make an initial assessment of whether population-level impacts are possible for the Northern Gannet, a species that has suffered significant global mortality, and two harvested species, Common Murre and American Common Eider, to support management decisions. We hypothesize that population-level impacts in eastern Canada are possible for Northern Gannets and American Common Eiders but are unlikely for Common Murres. This study underscores the urgent need for further research to understand the broader ecological ramifications of the HPAI outbreak on wild bird populations.

## Introduction

The Highly Pathogenic Avian Influenza A (HPAI) H5N1 virus of the A/Goose/Guandong/1/96 lineage emerged in Asia in 1996. Over the past 25 years, the virus has continued evolving and been detected across Asia, Africa, and Europe. There has also been an increasing number of mass mortality events involving wild birds (Verhagen et al. 2021), but until 2022, no unusual mass mortalities caused by HPAIV have occurred in North America (Papp et al. 2017, Giacinti et al. 2023). The current wave, clade 2.3.4.4b lineage H5NX HPAI with H5N1 as the dominant subtype, has marked a drastic turning point with wild birds becoming not only a reservoir, but also susceptible to morbidity and mortality associated with the virus. The HPAI virus is now considered the cause of the largest avian panzootic to date, based on the number of dead birds and species affected and the number and geographic spread of outbreaks (Klaassen and Wille 2023).

Globally, this new HPAI virus subtype has caused disease and unprecedented mass mortality events among wild birds with disproportionate impacts on waterbirds, including cranes (Lublin et al. 2023, Pawar et al. 2023), Great Skuas (*Stercorarius skua*; Banyard et al. 2022), gulls and terns (Pohlmann et al. 2023, Roberts et al. 2023, Sobolev et al. 2023), gannets (Lane et al. 2023, Pohlmann et al. 2023, Roberts et al. 2023), Peruvian pelicans (*Pelecanus thagus*; Leguia et al. 2023), Cape Cormorants (*Phalacrocorax capensis*), and African Penguins (*Spheniscus demersus*; Roberts et al. 2023). In North America, this virus was first detected in wild birds in December 2021, following observation of several wild gulls with neurological symptoms at a rehabilitation facility in St. John’s, Newfoundland, Canada (K. Gosse, pers. comm.). One of these, a Great Black-backed Gull (*Larus marinus*) was the index case for H5N1 2.3.4.4b in North America (Caliendo et al. 2022). During the following spring, mass mortalities involving thousands of wild birds were reported across eastern Canada, with the HPAI virus implicated as the cause.

Wildlife diseases that cause unusual levels of mortality or reduced fitness can exacerbate population declines when they interact with the cumulative effects of other natural and anthropogenic stressors which face marine birds (Phillips et al. 2023). Information on the scope and scale of unusual mass mortality events are needed to facilitate the assessment of population-level impacts, and support conservation and management decisions (e.g., species status assessments, harvest management), as well as being an important complement to pathogen surveillance programs that track the epidemiology of a disease and the evolution of the pathogen. However, robust assessments of mortality can be challenging to conduct, especially over vast areas. During large-scale mortality events, several types of surveys are employed to estimate the number of affected birds, along with monitoring the temporal and geographic scope of the emergency. For coastal species, beached bird surveys are typically conducted to estimate the number of birds affected by oiling and disease (e.g., Camphuysen 1998, Haney et al. 2014). Aerial (O’Hara et al. 2009) and boat-based surveys (Murphy et al. 1997) are also employed during environmental emergencies to estimate wildlife mortality and population impacts.

In this study, we provide the first comprehensive collation and assessment of wild bird mortality in eastern Canada during April - September 2022 following the fall 2021 incursion of H5N1 2.3.4.4b into eastern North America. Due to the unprecedented magnitude and geographic scale of the mortality, conducting systematic beached bird surveys to assess mortality was logistically infeasible. Instead, we combined data from federal, Indigenous, provincial, and municipal governments, the Canadian Wildlife Health Cooperative, non-governmental organizations, universities, citizen science platforms, and reports from the public to estimate the minimum number of birds of various species that died during this disease outbreak. In addition, we present an analytical approach for dealing with double-counted birds to address observations of mortalities which may have been reported by two or more different sources. With this corrected data set, we describe the magnitude of the unusual mass mortality event in terms of its spatial extent, duration, and the diversity and number of reported sick and dead birds. In addition to presenting mortality estimates, by comparing mortality numbers to population sizes and trends we make an initial assessment of whether population-level impacts are possible for three prioritized species: Northern Gannet (*Morus bassanus*, which suffered significant mortality globally (Lane et al. 2023), and two harvested species, American Common Eider (*Somateria mollissima dresseri*) and Common Murre (*Uria aalge*), to support harvest management decisions.

## Methods

### Study area and data sources

We defined eastern Canada as the provinces of Québec (QC), New Brunswick (NB), Nova Scotia (NS), Prince Edward Island (PEI), and Newfoundland and Labrador (NL), and we defined our study period as 01-April-2022 to 30-September-2022 to capture the period of greatest mortality. To generate the best available estimate of reported wild bird mortality linked with HPAI during the study period, we collated reports of sick and dead birds on land, on the water, and on breeding colonies. Information on recovery rates for wild birds is limited. We assume that birds reported as sick (i.e., with clinical signs consistent with HPAI infection including tremors, lack of coordination, or lack of energy or movement) succumb to infection (Roberts et al. 2023). Hereafter, both sick and dead birds are referred to as mortalities.

Reports of mortalities on land and water were collated from numerous sources including federal, provincial, Indigenous, and municipal government staff and databases, the Canadian Wildlife Health Cooperative and other NGOs, academic researchers, and two citizen science platforms (iNaturalist and eBird; see Appendix S1 for methods on extracting and processing mortality data). Observations of wild bird mortalities on seabird colonies visited by government biologists and academic researchers were obtained through direct solicitation. The reports included incidental observations made on any seabird colony, and standardized surveys that were conducted by boat, foot, or air. Unless reported mortalities were explicitly stated as having occurred on a colony, they were not classified as colony mortalities. All mortality data are provided in the online repository (Data S1).

Each record included the species, date, number of mortalities, location information (i.e., site name or coordinates), observer information (name and contact information), and information source. We anonymized observer information in the published dataset. When species assignments were not provided, less specific taxonomic assignments were used (e.g., unknown gull, unknown bird). Taxonomic identification for iNaturalist reports was verified where the quality of the species identification was rated as ‘needs_id’ by the submitter. When site names were provided instead of coordinates, coordinates were obtained using the GoogleMaps API in R version 4.2.2 (R Core Team 2023). A subset of coordinates and site names were reviewed by regional experts to confirm the validity of this approach. To provide broad estimates for species groups, we classified species as either: seabirds, waterfowl, waders, shorebirds, loons, landbirds, or raptors. Age or breeding status was recorded when available. To assess how age class may be differentially represented in the mortality events, photos of Northern Gannets submitted to iNaturalist (557) were reviewed and the age class of birds was classified as adult, subadult, hatch year (HY), or unidentifiable. Similarly, photos of Common Murres were reviewed (61) and birds were classified as HY, after hatch year (AHY), or unidentifiable. We assumed that the age structure of birds in these photos reflects the age structure of birds reported off colonies. Mortalities of adult birds on colonies were assumed to be breeding adults unless there was evidence to the contrary.

### Prioritized species

Environment and Climate Change Canada (ECCC), the federal wildlife management agency responsible for the conservation of migratory birds, prioritized three species based on a need for information to support conservation and harvest management decisions in 2022. Details about colony surveys in 2022 for Common Eiders, Northern Gannets, and Common Murres are summarized below and shared in the online data repository (Data S2, Data S3, Data S4, respectively).

#### Common Eiders

Eastern Canada is the core of the breeding range for American Common Eiders (Beuth et al. 2016, Lamb et al. 2019, Gutowsky et al. 2023), hosting 85% of the global breeding population in three subpopulations: QC North Shore and NL (60% of breeding birds), QC St. Lawrence Estuary (20% of breeding birds) and NB & NS (5% of breeding birds; C. Lepage, pers. comm.). The cryptic brown females of this species are wholly responsible for raising chicks, and nest in colonies on hundreds of islands across the region. In the St. Lawrence Estuary of QC, the largest breeding colony (Île Bicquette) has been declining for the last two decades (Lepage 2019). Other colonies in the St. Lawrence Estuary are stable or increasing (i.e., Île aux Pommes, Île Blanche, Île aux Fraises, Île aux Oeufs, Îles du Pot, and Île Laval; Giroux et al. 2021), as are the eider populations on the QC North Shore (Rail 2021b). Elsewhere in eastern Canada, populations have been declining (Giroux et al. 2021, Noel et al. 2021). Colonies in the St. Lawrence Estuary currently support recreational harvests in the USA and Canada (Rothe et al. 2015), and are the subject of down collection (Joint Working Group on the Management of the Common Eider 2004). An assessment of mortality and potential population impacts for this species was prioritized to support harvest management decisions.

To the extent possible, Common Eider colonies were surveyed by air or on foot throughout eastern Canada (Data S2). In QC, this included surveys on foot of the three largest colonies in the St. Lawrence Estuary, by down harvesters between May 29 and May 31 (Île Bicquette, Île aux Pommes, Île Blanche). Twelve colonies along the North Shore of the Gulf of St. Lawrence were surveyed on foot by ECCC, between May 29 and June 22, as part of the quinquennial colonial seabird monitoring program, which includes nine migratory bird sanctuaries. In NB, Machias Seal Island was partially surveyed weekly from mid-May to mid-August, and only one dead juvenile Common Eider was found. An additional six colonies were incidentally surveyed in the Wolves and the Grand Manan Archipelagos by ECCC staff between May 31 and June 5, 2022. Six colonies along the northern peninsula of the island of Newfoundland (NF) were partially or incidentally surveyed on foot and by boat by ECCC staff between May 14 and June 7. In Labrador, nine colonies along the south coast were incidentally surveyed by air (helicopter) and on foot between June 6 and July 12, 2022. For NS, two colonies (John’s Island and Grey’s Island in southwest NS) were incidentally surveyed by boat and on foot by ECCC staff between April 30 and May 1, 2022. Complete aerial (helicopter) surveys of 18 other NS colonies were conducted by the Nova Scotia Department of Natural Resources and Renewables staff on June 21, 2022. No known colonies exist in PEI.

#### Northern Gannets

Canada hosts the entire North American breeding population of Northern Gannets (213,704 breeding birds; Data: Table S2), which represented 13% of the global population as of the latest assessment (Mowbray 2020). Northern Gannets are long-lived seabird species that breed in dense colonies at three colonies in QC (Île Bonaventure, Rochers aux Oiseaux, Île d’Anticosti) and three colonies in NF (Funk Island, Baccalieu Island, Cape St. Mary’s Data: Table S2). Breeding populations at the five largest sites have increased dramatically since 1970 and are currently considered to be stable or increasing (i.e., from 2010-2020; d’Entremont et al. 2022, S. Wilhelm, pers. comm.).

In July and September 2022, ECCC flew aerial surveys of the two largest Northern Gannet colonies in QC (Île Bonaventure and Rochers aux Oiseaux) and all three Northern Gannet colonies in NF (Data S3). Aerial photographs were digitized following standardized procedures (Chardine et al. 2013, Rail et al. 2014). The number of Apparently Occupied Sites (AOS) and dead gannets were enumerated, in a complete survey of these colonies. Île d’Anticosti in QC has a very small population (192 breeding birds in 2019) and was not surveyed (J.-F. Rail, pers. comm.). In addition, complete surveys of 3 plots on Île Bonaventure were performed between July 11 and September 30, 2022. Both the colony at Île Bonaventure, and at Cape St. Mary’s are within provincial protected areas that are staffed during the summer months.

#### Common Murres

There are approximately 1.75 million Common Murre breeding in eastern Canada (Ainley et al. 2021). The majority breed in NF (1.5 million breeding birds), with smaller breeding populations in QC (191,500 breeding birds) and Labrador (76,228 breeding birds). The breeding population in NF and QC represent 19% of the North American population and ∼10% of the global population (Ainley et al., 2021). Common Murre breed in colonies at 68 sites in eastern Canada, including 39 sites in QC, 12 sites in NF, 15 in Labrador, and 2 sites in NB. Eight of these colonies are considered large (>20,000 breeding birds): three are in QC (Île Bonaventure, Rochers aux Oiseaux, Sainte-Marie Island), four are in NF (Funk Island, Green Island, Cape St Mary’s, South Cabot Island), and one is in Labrador (Gannet Islands). Common Murre are one of only two regulated harvested non-waterfowl marine species in Canada, and this species was prioritized for this assessment to support harvest management decisions.

In June 2022, the two small colonies at Machias Seal Island (NB) and Île à Calculot des Betchouanes (QC) colonies were completely surveyed on foot. An additional nine colonies were incidentally surveyed for evidence of mass mortality (Data S4). In July and September 2022, the colonies on Île Bonaventure, Rochers aux Oiseaux, Baccalieu Island and Funk Island were surveyed during aerial surveys of adjacent Northern Gannet colonies. Baccalieu Island was also incidentally surveyed on foot in August. In July, Gull Island and South Cabot Island were surveyed by boat and by helicopter, respectively. The colony at Cape St. Mary’s was surveyed by boat and helicopter in July and by boat in September. In July, Great Island and Gull Island in Witless Bay Ecological Reserve (NL) were incidentally surveyed by boat.

### Attributing mortality to HPAIV

A subset of sick and dead wild birds was tested for the HPAI H5N1 clade 2.3.4.4b virus in eastern Canada between April 1 and September 30, 2022, as part of Canada’s Interagency Surveillance Program for Avian Influenza Viruses in Wild Birds (*n* = 96 species; Giacinti et al. 2023). Given that our objective was to quantify mortality for the purpose of supporting evaluations of population-level effects and that no other notable sources of mortality were reported for this period (M. Jones, pers comm.), we assumed that HPAI was a likely cause of mortality for any species that tested positive for HPAI virus in sick and dead birds within our study region during our study period. For a description of the epidemiology of the HPAI outbreak in wild birds in Canada, including spatiotemporal dynamics, host taxonomic representation and viral genetic diversity see Giacinti et al., 2023.

For records where the species was unknown (e.g., unknown gulls), we presumed that mortalities were caused by the HPAI virus if ≥ 50% of the species within that group tested positive for the virus. For example, a bird identified as an “Unknown Gull” was likely one of any of the following species common in the study area: Herring Gull (*Larus argentatus*), Great Black-backed Gull, Ring-billed Gull (*Larus delawarensis*), Glaucous Gull (*Larus hyperboreus*), or Iceland Gull (*Larus glaucoides*). We checked if most of these species had tested positive for HPAI virus at some point within the study area and study period, and if yes, we presumed that mortalities of unknown gulls were linked to HPAI (Giacinti et al. 2023). We did not presume records identified only as ‘Unknown Bird’ were positive, and so mortalities for unknown birds were excluded from the HPAI mortality dataset, along with individuals that had a cause of death reported that could not be linked to HPAI and individual birds that tested negative for HPAI virus.

### Scenario analysis to identify double counted mortalities

Reported mortalities are subject to inflation when the same bird is reported by multiple observers, to multiple sources, or both (i.e., double counts). We presumed that when our dataset included records of a given species reported at similar times and places, this may indicate that an observation was captured in our dataset more than once. To identify records in the HPAI mortality dataset that could be considered observations of the same birds, we undertook a comparative analysis to examine how the number of reported mortalities, that could be considered double-counted, would change under six scenarios that explore a range of spatial and temporal overlap (Table 1).

**Table 1.** Total number of HPAI-linked mortalities reported (i.e., baseline scenario) and the estimated number of mortalities with double counts removed according to a range of criteria. Reported mortalities for birds with less specific taxonomic identities that were presumed positive but that could not be included in the double count analysis have been added to totals. The values do not include birds that tested negative for HPAI virus and species that were presumed negative or the 2,762 Unknown Birds, because these mortalities could not be attributed to the HPAI virus.

| Criteria | Baseline | Scenario A | Scenario B* | Scenario C | Scenario D | Scenario E | Scenario F |
| --- | --- | --- | --- | --- | --- | --- | --- |
| **Days** | N/A | ±0 day | ±1 day | ±5 day | ±0 day | ±5 day | ±1 day |
| **Distance** | N/A | ±1 km | ±1 km | ±1 km | ±5 km | ±5 km | ±5 km |
| **Total mortality** | 44,721 | 42,294 | 40,966 | 39,743 | 41,408 | 39,176 | 40,122 |
| **Δ from Baseline** |  | 2,427 | 3,755 | 4,978 | 3,313 | 5,545 | 4,599 |
| **%Δ from Baseline** |  | 5.4% | 8.4% | 11.1% | 7.4% | 12.4% | 10.3% |
| **Including:** |  |  |  |  |  |  |  |
| **Common Eider** | 1,983 | 1,971 | 1,945 | 1,912 | 1,961 | 1,926 | 1,940 |
| **Northern Gannet** | 28,406 | 26,842 | 26,193 | 25,653 | 26,320 | 25,249 | 25,541 |
| **Common Murre** | 9,092 | 8,716 | 8,133 | 7,652 | 8,433 | 7,552 | 8,103 |
| **Gull** | 2,378 | 2,348 | 2,326 | 2,260 | 2,330 | 2,181 | 2,260 |
| **Cormorant** | 997 | 990 | 981 | 959 | 988 | 948 | 950 |
| **Atlantic Puffin** | 364 | 299 | 282 | 244 | 259 | 276 | 238 |
| **Black-legged Kittiwake** | 260 | 257 | 251 | 227 | 252 | 224 | 245 |
| **Razorbill** | 127 | 121 | 119 | 109 | 120 | 107 | 118 |
| **Tern** | 83 | 80 | 74 | 75 | 80 | 74 | 74 |

\*Scenario B (±1 km, ±1 day) was selected as the best compromise between removing multiple records of the same birds and but not removing records of new batches of birds arriving in an area.

Specifically, the double count scenario analysis removes records from the complete mortality dataset as a function of the number of days between observations (±0 days, ±1 day, ±5 days), and the distance between observations (±1km, ±5km). The baseline scenario is the total reported HPAI-linked mortality with no records excluded and assumes no double counting. More precisely, each mortality record *a_i,j_* was coupled with another mortality record *a_i ≠ i,j_*, where *i* was a unique identifier for the record and *j* was the species. The distances (km) and the number of days between paired records were calculated and if the scenario conditions were met (e.g., for Scenario A, record *a_i,j_* fell within 1 km and 0 days of record *a_i ≠ i,j_*), the records were considered to be double counts. For each pair of records considered double counts, the record with the larger number of total observed birds was retained, and the other record was excluded. All retained records were iteratively re-submitted for consideration (i.e., coupled with another record within the dataset) until all records that met the scenario conditions were considered, and only records that did not meet the scenario conditions (i.e., non-double counts) and double counts with the highest number of total observed birds remained. Both records in a pair were retained only if the reports were made by the same observer to the same source. This exception reduced the mistake of excluding reports made in close proximity, as is typical during beached bird surveys.

We assumed taxonomic identifications were accurate, and reports were only considered as possible double counts if the species’ name matched. However, we acknowledge that some observers may have mistakenly identified one species as another (e.g., a Great Black-backed Gull for a Herring Gull). To address this possibility in the double count analysis, we treated reports of cormorants, gulls, and terns as interchangeable regardless of the species or taxonomic resolution (Appendix S2: Table S1). For example, paired reports of Arctic Tern (*Sterna paradisaea*), Common Tern (*Sterna hirundo*), and Unknown Tern were considered potential double counts if they met the criteria. We did not perform the double count analysis on reports that had other non-specific taxonomic assignments (e.g., unknown alcid) because it was not reasonable to consider them to be interchangeable.

To arrive at a double-count corrected estimate of total reported HPAI-linked mortality for each scenario, we discarded records identified as double counts and added all recorded mortalities for species with less-specific taxonomic status (except for the cormorants, gulls, and terns) if those unknown species were presumed to be positive for HPAI. We compared the total reported HPAI-linked mortality from the various scenarios to the total reported HPAI-linked mortality with no records excluded. To present spatiotemporal patterns and species-specific information about mortality, we used a double-count corrected estimate. based on consultation with experts who have experience with similar datasets and research questions. to choose the scenario which best represented a reasonable compromise between excluding double counts and retaining unique records.

#### Species-specific mortality estimates

We calculated species-specific mortality estimates after removing records unrelated to HPAI and removing double counts based on the chosen scenario. Mortality estimates were calculated by summing all records for a particular species. For the three prioritized species (Common Eider, Northern Gannet, Common Murre), we present a more comprehensive spatiotemporal analysis of mortality events in the study area, with specific dates and events highlighted. The summed mortality numbers presented are minimum estimates, as we did not correct for birds that died that went unrecorded (e.g., areas not surveyed, birds not detected, birds not reported, birds lost at sea).

## Results

### Reported mortalities

In total, 48,042 wild bird mortalities were reported in eastern Canada between April 1 and September 30, 2022 (Data S1). This complete dataset of reported mortality includes 142 species, as well as 23 less specific taxonomic assignments. Most of the mortality data were provided by governmental bodies. Notably federal staff from ECCC (28.3%), the Provincial wildlife management and natural resources departments of QC, NS, NB, PEI, and NL (18.6%), and municipal governments (10.8%) contributed to a substantial number of the reported mortalities. Mortality data were also reported by Parks Canada (2.0%), Fisheries and Oceans Canada (1.8%), and Indigenous governments in Labrador (Nunatsiavut Government, NunatuKavut Community Council) and the island of Newfoundland (Miawpukek First Nation and Qalipu First Nation) (0.04%). Non-governmental organizations, including the Canadian Wildlife Health Cooperative, contributed 7.7% of the reported mortalities, while academia provided 17.6%. The reports of citizen scientists made to iNaturalist and eBird contributed 0.9% and 0.8% of reported mortalities, respectively. The remaining 11.5% of reported mortalities were made by members of the public directly to ECCC.

### Scenario analysis

After removing records for 3,321 mortalities that could not be attributed to HPAI, our baseline HPAI mortality dataset includes 44,721 wild bird mortalities (Table 1). Across the scenarios, 5.4% to 12.4% of reported mortalities were identified as potential double counts (Table 1). In the most restrictive scenario (Scenario A, ±0 day, ±1 km), 2,427 birds were identified as potential double-counts, and 5,545 birds were identified in the least restrictive scenario (Scenario E, ± 5 day, ± 5 km; Table 1). A comparison of scenario results indicates that the closeness in time criteria had a larger influence on the number of birds identified as double counts than the distance criteria. The two scenarios that used the ±5 day criteria (i.e., Scenarios C and E) identified the highest percentage of birds as double counts (11.1%, and 12.4%, respectively). The two scenarios that used the ± 0 day criteria (i.e., Scenario A and D) identified the smallest percentage of birds as double counts (5.4% and 7.4%, respectively). Scenarios B (± 1 day, ± 1 km) and F (±1 day, ± 5 km) were closest to the median percentage of records identified as double counts.

We felt that Scenario B most likely represents the true reported mortality (Table 1) because it balances accounting for introduced errors associated with our georeferencing of site names and not excluding new birds that may come ashore each day. Values presented in the remainder of the paper reflect Scenario B results. In Scenario B, most records identified as double counts occurred in NL (51.4%), NS (28.1%), and QC (15.7%), with smaller double counts occurring in the other provinces (Appendix S2: Figure S1). Most double counts were reports of Northern Gannets (58.9% of reports, representing 2,213 birds) and Common Murres (25.5% of reports, representing 959 birds).

### Spatial, temporal, and taxonomic patterns of mortality

Accounting for double counts we estimate 40,966 individual birds of 45 species were reported in eastern Canada between April 1 and September 30, 2022 (i.e., Scenario B; ±1 km, ±1 day, Table 1). The first large wave started in May when mass mortalities were first detected among Common Eiders in the St. Lawrence Estuary, followed by mass mortalities of Northern Gannets and other species (e.g., gulls, cormorants, alcids) in the southern Gulf of St. Lawrence. This mortality event continued until mid-September. A second large wave of mortality began in June in eastern Newfoundland, with reports of Northern Gannets, Common Murres and other species starting in the southeast (e.g., Burin and Avalon Peninsulas) and later along the eastern and northeastern Newfoundland coasts. Mortalities continued to be reported until mid-September. The high numbers of colony mortalities for Northern Gannets reported the week of July 11 and September 12 reflect a reporting lag, as periodic aerial surveys of colonies provide a snapshot of mortalities during prior weeks. Reports of mortality along QC’s North Shore, southwestern Newfoundland, and in Labrador were limited.

Seabirds and sea ducks accounted for 98.7% (40,438) of reported mortality, with much smaller numbers of waterfowl (282, 0.7%), landbirds (133, 0.3%), raptors (97, 0.2%), and waders (16, 0.04%). Among seabirds, Northern Gannets accounted for 63.9% of all reported mortality (26,193). A significant proportion of reported mortalities were Common Murres (8,133, 19.9%), with smaller numbers of gulls (2,326, 5.7%), Common Eiders (1,945, 4.8%), cormorants (981, 2.4%), Atlantic Puffins (*Fratercula arctica*, 282, 0.7%), Black-legged Kittiwakes (*Rissa tridactyla*, 251, 0.6%), Razorbills (*Alca torda*, 119, 0.3%), and terns reported (74, 0.2%). Among waterfowl, land birds, raptors and waders, the most reported species were Canada Goose (*Branta canadensis*, 107, 0.3%), American Crow (*Corvus brachyrhynchos*, 99, 0.2%), Bald Eagle (*Haliaeetus leucocephalus*, 43, 0.1%), and Great Blue Heron (*Ardea herodias*, 15, 0.04%).

Approximately 45% of reported bird mortalities were observed on seabird colonies, and the remaining 55% of mortalities were reported elsewhere, on land and water. On-colony mortalities were dominated by Northern Gannets in QC (9,177) and the island of Newfoundland (4,899), with smaller mortalities of other species being reported on colonies in NB (963) and NS (643; Figure 1B). The largest numbers of bird mortalities were reported on land and water in NL (8,738 birds, including 100 birds in Labrador), although significant mortalities were also reported in QC (7,076), NB (4,461), and NS (1,522). Fewer than one thousand wild bird mortalities were reported on PEI (915).

**Figure 1.**
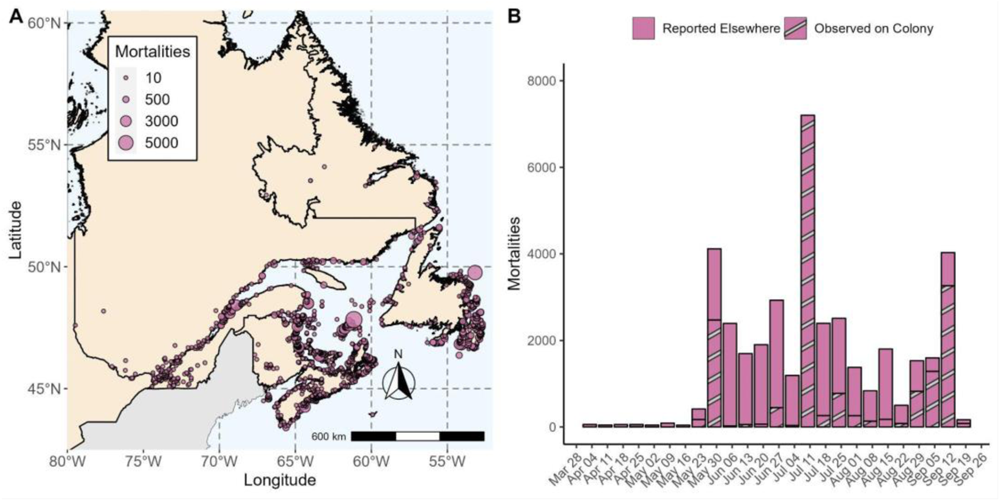
Spatial (A) and temporal (B) distributions of HPAI-linked wild bird mortality in eastern Canada, April 1 – September 30, 2022. These values represent the estimates derived from the double count analysis (Scenario B: ±1 km, ±1 day).

### Summaries for prioritized species

Accounting for double counts (i.e., Scenarios B (± 1 day, ± 1 km) and F (±1 day, ± 5 km)), Northern Gannets, Common Murres, and Common Eiders suffered mass mortality during the HPAI outbreak, and we provide an overview of the spatiotemporal distribution of the mortalities for these species. Similar information for gulls, cormorants, Atlantic Puffins, Black-legged Kittiwakes, Razorbills, and terns is provided in Appendix S3.

#### Common Eiders

In total, 1,945 Common Eider mortalities were reported in eastern Canada during our study period (Figure 2A). Most were breeding females observed at three colonies in the St. Lawrence Estuary: Île Bicquette (610), Île aux Pommes (503) and Île Blanche (222). These mortalities were reported during regular down harvesting operations (when complete surveys of the colonies are carried out) in May and beginning of June, and as result of follow-up partial surveys at the end of June.

**Figure 2.**
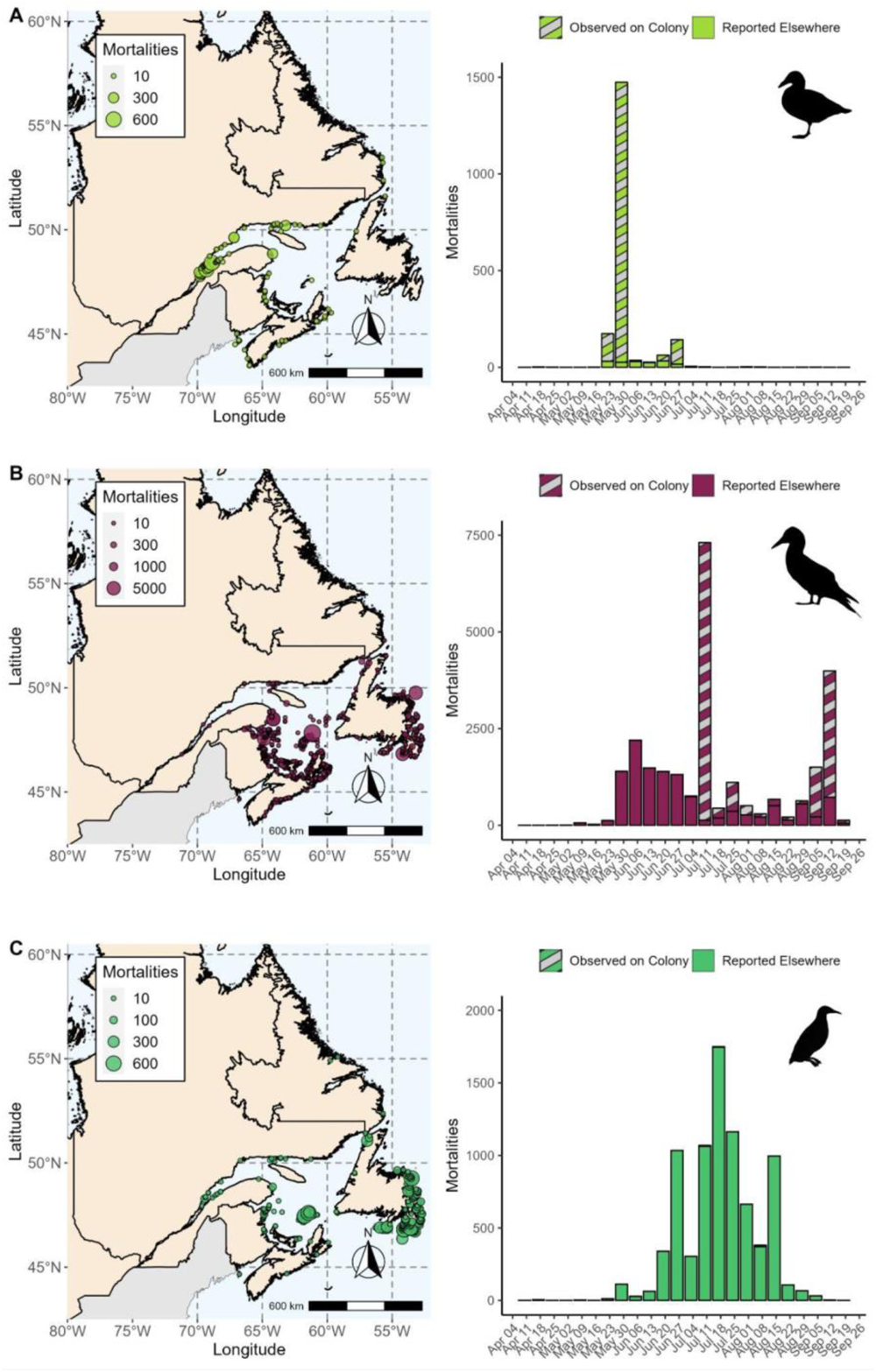
The spatial and temporal distribution of HPAI-linked mortalities across eastern Canada for (A) Common Eider, (B) Northern Gannet, and (C) Common Murre, reported between April 1 and 30 September 2022, with double counted birds removed (Scenario B: ±1 km, ±1 day).

HPAI-linked mortality was also reported at several other smaller Common Eider colonies in the St. Lawrence Estuary: Île aux Fraises (66), Île Le Gros Pot (44), and Île Le Pot du Phare (3). Unusual mortality was also reported at Île La Razade d’en Bas (67), Île La Razade d’en Haut (10), Île aux Lièvres (53), and Île aux Basques (30). Other known eider breeding colonies in QC, NB, and NL were surveyed in May and June with no unusual mortality being detected. In June, known breeding islands in NS were visited and no dead eiders were observed. Few HPAI-linked mortalities of eiders were reported off-colonies in eastern Canada (139 in QC, 21 in NS, 13 in NB, 5 in NL (of which 2 were on the island and 3 in Labrador) and none reported in PEI).

#### Northern Gannets

During our study period, 26,193 Northern Gannets mortalities were reported in eastern Canada (Figure 2B). Approximately half the mortalities were reported on the five gannet colonies that were surveyed (14,091), and half were reported on beaches and on the water (12,102). The mortality occurred in two waves: the first was reported in the Gulf of St. Lawrence in QC in May and involved at least 19,695 gannets. Approximately a month later, in June, mass mortality was detected in eastern Newfoundland involving at least 6,498 gannets. Both mortality events lasted until mid-September.

Out of 557 images of individual dead gannets submitted with iNaturalist reports, 92.8% were identified as adults. Another 3.2% were categorized as subadults based on the presence of black secondaries or tail feathers and it was not possible to identify the age of birds in the remaining 4% of images. Consequently, we assume that 11,231 out of 12,102 dead gannets reported off colonies were adults, and many would be expected to be breeding individuals.

The dominant source of the first wave of mortality is inferred to be the Rochers aux Oiseaux colony north of Îles-de-la-Madeleine in QC, based on the large number of carcasses observed on nearby beaches (4,065). Rochers aux Oiseaux is a remote colony in the middle of the Gulf of St. Lawrence and hosts 25% of Canada’s population of breeding gannets (∼52,000 breeding birds; Mowbray 2020, Rail 2021a). The earliest documentation of an outbreak and mass mortality at this colony was from Transport Canada’s National Aerial Surveillance Program on June 27, and subsequent aerial population surveys on July 11 detected a total of 5,119 dead adults. Dead gannets were not counted during the September aerial survey to avoid double counts, as it was difficult to distinguish between old and new carcasses.

To the west, at the Île Bonaventure colony, which hosts 50% of Canada’s breeding gannets (∼104,000 breeding birds; Rail 2021a), mortality was also reported (3,609). Île Bonaventure is closely monitored and the first on-colony report of mortality was made on May 24, 2022, by an employee of Parc National de l’Île-Bonaventure-et-du-Rocher-Percé (J.-L. Legault, pers. comm.). A subset of these birds tested positive for HPAI virus (Giacinti et al. 2023). In addition to quantifying dead birds observed in the July 11 aerial population surveys, dead birds in three study plots were enumerated weekly between June 18 and October 10. The mortality rate in study plots peaked between July 29 and September 10, but continued until the Park closed October 10, 2022 (Seyer et al., in prep.). Only mortalities reported in plots after July 11 are included in the mortality tally for this colony, to avoid double counting carcasses enumerated from aerial photographs. Direct counting of dead gannets indicated a smaller proportion of the Île Bonaventure breeding population died of HPAI (minimum of 3.5% colony mortality) than at Rochers aux Oiseaux (minimum of 9.8% colony mortality), although effort was unequal and the complete colony survey at Île Bonaventure preceded the peak of the mortality event.

The second wave of mortalities reports started in southeastern Newfoundland in early June and lasted until the second week of September. Outbreaks were detected at all three colonies in this region, which together host 25% of Canada’s population of breeding gannets (Cape St. Mary’s ∼30,000 breeding birds; Funk Island ∼22,000 breeding birds; Baccalieu Island ∼7,000 breeding birds; S. Wilhelm et al., in prep.). The first mortality to test positive for HPAI near a colony was a subadult found by a fisherman within 1km of Cape St. Mary’s on June 5, 2022 (E. White, pers. comm.). The first mortalities among breeding birds were first recorded by employees at Cape St. Mary’s Ecological Reserve on July 16^th^, 2022 (C. Mooney, pers. comm.). Additional mortalities were observed at this site until the onset of fledging on September 9. Population surveys at Cape St. Mary’s colony detected 1,551 mortalities (at least 5.2% colony mortality) and 1,050 dead birds reported on the southeastern coastline of Newfoundland likely originated from this population.

The colonies at Funk and Baccalieu are remote, and the status of outbreaks was only evaluated during aerial surveys. The first aerial surveys of the Funk and Baccalieu colonies on July 27 and July 24, respectively found only a limited number (149, 13 respectively) of dead individuals on these colonies; however later flights in September (September 15, Funk and September 14, Baccalieu) revealed a much larger-scale die-off. This suggests an outbreak occurred some time between late July and mid-September. At the Funk Island colony, 3,158 mortalities were observed (at least 14.4% colony mortality), while at the smaller Baccalieu Island colony, 28 mortalities were observed (at least 0.4% colony mortality). The 545 birds reported along the northeast and eastern shores of Newfoundland likely originated from these colonies.

#### Common Murres

Across eastern Canada, 8,133 Common Murre mortalities were reported (Figure 2C). The mortality event for Common Murres followed generally the same pattern as Northern Gannets, with HPAI-linked mortality being reported first in the Gulf of St. Lawrence in QC (2,422), followed by an outbreak in eastern Newfoundland (5,708). The larger mortality event on the island of Newfoundland is consistent with the larger population (∼1.5 million breeding birds) compared to the Gulf of St. Lawrence (191,500 breeding birds; Ainley et al. 2021). Most of the 61 images of Common Murres associated with iNaturalist mortality reports were AHY (86.9%). Only 3.3% were HY birds (i.e., post-fledge chicks), and 9.8% of images were unidentifiable. Consequently, we assume that 7,049 of the 8,112 dead Common Murres reported off colonies were adults (86.9%) and may have been breeding individuals at a minimum.

In the Gulf of St. Lawrence, mortalities started in late May and lasted until mid-August, with most being reported between June 20 and July 20 on beaches of Îles-de-la-Madeleine.

These birds likely originated from colonies in Îles-de-la-Madeleine including Rochers aux Oiseaux, which host ∼54,000 breeding murres (Ainley et al. 2021). Smaller numbers were reported on the Gaspé Peninsula and in northern NB, near a large murre colony at Île Bonaventure (∼80,000 breeding murres; Ainley et al. 2021). Although an estimated ∼50,000 murres breed at colonies in this area, no mortalities were observed at either murre colony and few murres were reported on the QC North Shore of the Gulf of St. Lawrence.

A second, larger wave of reported mortalities event started approximately a month later, in late June, on the island of Newfoundland. Initial reports of mortalities came from the Burin Peninsula and Avalon Peninsula in southeastern Newfoundland. These birds likely originated from the colonies in the Cape St. Mary’s and Witless Bay Ecological Reserves, which together host ∼540,000 breeding murres (Ainley et al. 2021). By August 2022, the outbreak progressed northwards, with birds reported on the Bonavista Peninsula and Bonavista Bay. At least some of these birds likely originated from South Cabot Island (∼ 20,000 breeding birds), based on the recovery of at least one bird off the northwest coast of Bonavista Bay that was banded as a chick on South Cabot Island in 2018. This bird was recovered dead on Ship Island (49.0642, -53.5693) on 4 August 2022, and subsequently tested positive for the HPAI virus (Giacinti et al. 2023). Few murre mortalities were reported in southern Labrador (7).

No mass mortality was observed at any of the Common Murre colonies that were surveyed. Only 0.3% of reported mortalities were in colonies (21). This includes small numbers of mortalities (≤ 10) observed at Cape St Mary’s, and the Great and Gull Island colonies in the Witless Bay Seabird Ecological Reserve

## Discussion

The HPAI H5N1 2.3.4.4b virus is now considered the cause of the largest avian panzootic to date, based on the number of dead birds, species affected, and the number and geographic spread of outbreaks (Klaassen and Wille 2023). Our study provides the first comprehensive assessment of reported wild bird mortalities in eastern Canada during the mass mortality caused by the HPAI H5N1 2.3.4.4b virus between April and September of 2022. To establish a minimum estimate, we collated data from multiple sources in a comprehensive effort to understand regional spatiotemporal and taxonomic patterns of mortality both on and off seabird colonies. We presented a scenario-based method for identifying and excluding observations that were potentially reported more than once by different observers or sources (i.e. double counted).

After the double-counts analysis, we conclude that the HPAI virus is linked to the deaths of at least 40,966 wild birds between April 1 and September 30, 2022. Mass mortality was restricted to the eastern Canadian provinces of QC, NB, PEI, NS, and insular NF. Limited mortality was reported in Labrador. Experts indicated no wild bird mortalities on this scale (i.e., in the thousands) were reported during the study period on the eastern coast of the USA (Atlantic Marine Bird Cooperative, Community Science and Marine Bird Health Working Group, pers. comm.). To our knowledge, in North America, there has been no recorded infectious disease that has caused a comparable level of mortality across such a diverse range of bird species. The mortality reported here far exceeds mortalities associated with the 2014-15 outbreak of HPAI in waterfowl in the prairies (Canadian Food Inspection Agency 2016a, 2016b), or avian cholera outbreaks in prairie waterfowl (1977; Wobster et al. 1979), Arctic seaduck colonies (mid-2000s; Iverson et al. 2016), or Alaskan seabirds (2013; Bodenstein et al. 2015), although larger mortality events caused by marine heat waves have been reported (e.g., Piatt et al. 2020). In our study, we found that between 5.4% and 12.4% of reported mortalities may have been reported by more than one individual or source, highlighting the importance of accounting for double-counts during large scale data collation exercises. Our approach for handling double-counted mortalities could be applied to support the assessment of mortality in any case where information from multiple sources are used.

In our study, Northern Gannets, Common Murres and Common Eiders suffered the greatest mortalities, followed by gulls, cormorants, Atlantic Puffins, Black-Legged Kittiwakes, Razorbills, and terns. Importantly, mortality estimates from scenarios that remove possible double counts may be more accurate than the baseline scenario, but they still fall short of capturing the complete picture because not all dead birds are reported. Below, we discuss the potential for population-level impacts for the three prioritized species: Common Eiders, Northern Gannets and Common Murres. Given the size of the breeding populations for these species in eastern Canada relative to the number of mortalities reported, population-level impacts are certainly possible and even likely for Northern Gannets throughout their whole Canadian range, and for regional populations of Common Eiders, but are not likely for Common Murres.

### Common Eiders

A notable event occurred in the St. Lawrence Estuary in QC, where mass mortalities were observed, affecting the colonies at Île aux Pommes, Île Bicquette, and Île Blanche. These three colonies represent 58% of the breeding population in the St. Lawrence Estuary and 11% of the breeding population of Common Eiders in Canada, and the USA (Lepage 2019; C. Lepage, pers. comm.). When compared their colony size in the previous summer (2021; 4619, 3263, and 3155, breeding pairs (C. Lepage pers. comm.)), the HPAI outbreak resulted in 13.2%, 16.1%, and 6.8% mortality, respectively. Cumulatively, the reported mortality at these three islands represents a 12.1% loss of breeding females from these three sites during early incubation. As a result of the mortality event, we expected the number of young birds produced to have been particularly low in 2022 and the population to be lower in the coming years.

Given that breeding colonies in the St. Lawrence Estuary support recreational harvests and the collection of eider down, this species was prioritized to support harvest management decisions. After considering the potential population level impacts of HPAI-linked mortality, population trends, and harvest pressures, ECCC’s Canadian Wildlife Service – the agency responsible for hunting regulations in Canada under the Migratory Bird Act - recommended *“a voluntary reduction in eider harvest for the 2022-2023 season and that hunters refrain from harvesting female common eider or young”* (Environment and Climate Change Canada 2022). Changes to the migratory bird hunting regulations, to reduce the impact of the harvest on the Common Eider population in QC, was not possible given the short period of time between the assessment of the mortalities on colonies and the date of the hunting season opening. Therefore, awareness and cooperation of hunters was deemed likely sufficient to mitigate the risk of an excessive decline in this population. ECCC also reached out to the different U.S. States agencies that manage Common Eider hunting (e.g., Maine Department of Inland Fisheries & Wildlife, Massachusetts Division of Fisheries & Wildlife) to relay this message about voluntary restriction to local hunters (C. Lepage, pers. comm.). Ongoing annual nest monitoring at eider colonies in the St. Lawrence Estuary will support the assessment of the long-term impacts of HPAI mortality on these populations.

Outside of the St. Lawrence Estuary, no other major mortality events were reported in colonies across the North Shore of QC, in NS, NB, and NL. Although these colonies were only incidentally surveyed and fewer than 178 dead Common Eiders were reported off colonies across eastern Canada, we are reasonably confident that no mass mortality events occurred outside the St. Lawrence Estuary. This is fortunate because the NB and NS subpopulation is declining for reasons that are not fully understood (Milton et al. 2016, Giroux et al. 2021, Noel et al. 2021). It is worth noting that Common Eider mortalities were also reported in Maine (10), Massachusetts (19), and New Hampshire (1), and subsequently tested positive, between June 24, 2022, and July 25, 2022, (USDA 2023) but no mass mortality events at colonies have been reported (C. Lepage, pers. comm.).

### Northern Gannets

HPAI-linked mortality was observed at all five of the surveyed colonies. The second and fourth largest colonies, Rocher aux Oiseaux and Funk Island, appear to have suffered massive outbreaks, while the outbreak at Île Bonaventure was less dramatic but may have lasted longer. As large white birds with well-defined territories, gannets are among the easiest to enumerate from aerial surveys (Chardine et al. 2013), and yet it was still challenging to differentiate between old and new carcasses during the two aerial surveys of QC gannet colonies, where the main mortality event occurred prior to the first aerial survey in July. For this reason, mortality at the QC colonies was only estimated from the July photographs and should be viewed as minimums. On NF, the main mortality event occurred after the first survey and before the second survey, so mortalities were enumerated in aerial photographs from both surveys.

We can assume that the vast majority of Northern Gannets that were observed dead on breeding colonies (14,091) were adults, and that 12,102 of the mortalities reported off colonies were also adults (i.e., 92.8%). If those adults were breeding individuals, then the reported mortality represents a minimum 11.6% loss of the North American breeding population. However, this should be viewed as an absolute minimum. Many Northern Gannets were likely lost at sea (e.g., Polhmann et al., 2023), particularly those that died in the waters off the northeastern coast of Newfoundland (i.e., Funk Island, Baccalieu Island), where the dominant currents (Wu and Tang, 2011) would tend to advect birds away from shore. Efforts to estimate at sea losses and population surveys to detect changes in Apparently Occupied Territories (AOTs) will help to resolve the complete picture of population impacts from the HPAI virus. For example, while the number of mortalities enumerated at Rocher aux Oiseaux represents only 9.8% of the breeding population, a 58% decline in the number of Apparently Occupied Territories was observed in 2022, compared to the previous survey in 2020 (J.-F. Rail, pers. comm.).

Since monitoring of Northern Gannet colonies began in 1970, populations have increased significantly (Chardine et al. 2013). Across all five main colonies, the North American population is increasing or stable (Rail 2021a, d’Entremont et al. 2022, S. Wilhelm, pers. comm.). A full assessment of the population-level impact of HPAI on the six Northern Gannet populations in eastern Canada, and how long populations will take to recover, is beyond the scope of this paper. Several factors will influence recovery, including the number of mature non-breeding individuals available to recruit into the breeding population, breeding success in subsequent years, acquired immunity to future infection, and the cumulative impacts of natural and anthropogenic stressors that may have additive or synergistic effects with HPAI. Epidemiology, including testing for active infections and serology in subsequent years, will be important to understand exposure, survival, and immunity (Giacinti et al. 2023).

Globally, Northern Gannets are listed as Least Concern because they have a very large range and a large population size (1,500,000 to 1,800,000 mature individuals; BirdLife International 2023). However, Northern Gannets were impacted across their range, with an estimated 75% of colonies experiencing unusual mortality events (Lane et al. 2023). This includes marked declines in the number of breeding birds at key colonies in the UK (e.g., Bass Rocks, over 71% decline in June 2022 compared to 2014; Lane et al. 2023). To fully appreciate the population-level impact of HPAI on gannets throughout their range, an analysis of changes in breeding populations before and after mortality events will be needed. Fortunately, breeding Northern Gannets have well-defined territories and are methodologically easy to census via aerial surveys, and innovative approaches to cost-effectively survey populations are in development (Kuru et al. 2023 Walker et al., in prep).

### Common Murres

In eastern Canada, Common Murres were the second most reported species, with 8,133 Common Murre mortalities. We estimate that 7,049 of these were adults and may have been breeding. This mortality represents less than 0.5% of the breeding populations, therefore clear population-level impacts of HPAI-linked mortality in 2022 is not expected. However, the dominant currents near the largest murre colonies on the eastern and northeastern coasts of NF likely advect birds away from shore (Wu and Tang 2011). If at-sea losses are significant, the total number of murres succumbing to HPAIV may be severaly underestimated based on reported mortalities. As with gannets, efforts to estimate at-sea losses in 2022 and population surveys at key colonies (i.e., those with large populations) in the years post-outbreak will support a better understanding of population level impacts for Common Murres.

Unlike Northern Gannets, no mass mortalities of Common Murre were observed directly on breeding colonies in eastern Canada. The lack of mortalities seen on colonies themselves was surprising given that thousand of adult murre mortalities were reported on nearby beaches. Similar observations have been made in Europe (Germany, E. Ballstaedt pers. comm). Initially, we reasoned that murre mortalities may not be readily observed at the colony because most murres nest in packed colonies on rocky cliff ledges, and sick or dead birds had fallen into the water. However, on Funk Island and Rocher aux Oiseaux, murre and gannets breed in adjacent colonies on the plateau, but still no dead murres were observed. We speculate that murres leave the colony when sick and recommend that characterizing the magnitude of outbreaks among Common Murre based on mortality observed on the colony may not be the best approach.

Common Murre are harvested in Newfoundland and Labrador, a culturally important practice in the province (Chardine et al. 2008). This species is numerous in eastern Canada and populations have been increasing in QC, increasing or stable in the island of Newfoundland, and declining in Labrador (Ainley et al. 2021). After reviewing abundance, population trends and the mortality data presented here, ECCC chose “*not to change migratory bird hunting regulations to reduce the harvest of Murres in Newfoundland and Labrador during the 2022 to 2023 hunting season”* (Environment and Climate Change Canda 2022).

### Considerations for Interpreting Reported Mortalities on Beaches and at Sea

It is important to acknowledge two simplifications that were made in arriving at these reported mortality estimates. First, we attributed all mortalities to HPAI for species that tested positive for the HPAI virus in the study area during the study period, removing only those records where individual birds tested negative. Species specific HPAI virus prevalence rates are available from Canada’s Interagency Surveillance Program for Avian Influenza (Giacinti et al. 2023), but using these to adjust mortality estimates yielded unlikely results. For example, in eastern Canada, prevalence of HPAI virus in Northern Gannets was 62.8%. If we attributed only 62.8% of reported mortality to HPAI, this would suggest that 9,744 adult Northern gannets died of unknown causes. This is far above background annual mortality rates for adults of this species in North America (5% per annum; Chardine et al. 2013). Attributing all mortalities to the HPAI for species that tested positive for the virus in the study area during the study period may be simplistic, but in the absence of evidence that another large-scale morality factor is operating at the same time it seems a reasonable assumption. It is it is also consistent with other studies assessing HPAI mortality (e.g., Rijks et al. 2022, Pohlmann et al. 2023).

The second simplification was the assumption that all sick birds die. This was necessary, as the status of birds (sick or dead) was not consistently reported. While some birds may recover from HPAI (Lane et al. 2023), even when they exhibit severe neurological symptoms (L. Taylor, pers. comm.), most do not. The assumption that all sick birds died would only cause an overestimate of reported mortality if a large proportion of sick birds recovered. To our knowledge, recovery rates of seabirds’ with HPAI are low (e.g., African Penguins; Roberts et al. 2023). Although these simplifications could lead to an over-attribution of reported mortalities to HPAI, we are confident that our estimates still represent only a fraction of the total mortality resulting from the HPAI virus outbreak in eastern Canada.

There are several reasons we view reported mortalities as a minimum estimate. Our estimates are based on reported observations of sick and dead birds across an area spanning 5 Canadian provinces and Atlantic coastlines across 15 degrees of latitude. Although reporting efforts were significant it is likely that many bird carcasses were not observed or reported, even in areas of intensive effort. A considerable body of literature shows that detection rates and persistence probabilities for seabird carcasses on beaches is higher for large birds and white birds than it is for small birds and dark birds, and that beach type, weather, and degree of scavenging all influence detection and persistence (e.g., Fowler and Flint 1997, Wiese and Robertson 2004). It was beyond the scope of this study to generate an extrapolated number to estimate total carcasses based on these factors (i.e., grounding probability, detection probability, persistence rate) due to the sheer size and geographic scope of this outbreak.

Additionally, detection and reporting of mortalities is likely to be lower in areas with lower human population density. Preliminary drift modelling suggests that large numbers of gannet carcasses likely deposited on the North Shore of QC and on the shores of Île d’Anticosti (Avery-Gomm et al. 2023), but very few mortalities were reported in these areas, which are very remote from large population centers. Therefore, low reported mortalities in remote areas may reflect low detection due to low human population density rather than low mortalities. Finally, not all reported observations were captured due to capacity challenges of some jurisdictions to receive and record large volumes of calls from the public during the summer months when other urgent priorities arose (e.g., wildfires).

In eastern Canada, almost all reported mortalities were among seabird and seaduck species that forage at sea. Among such species, sick individuals may have died at sea and at-sea losses of these carcasses may comprise a significant fraction of total mortality. Experiments designed to assess at-sea loss in the context of large oil spills and chronic oil pollution have identified that temperature, scavenging activity, and body size are some factors that influence how long a carcass will float (Burger 1991, Ford et al. 1996, Wiese 2003). Carcasses that float for longer have a higher probability of washing ashore (i.e., grounding), but grounding probabilities are strongly influenced by oceanographic conditions, wind (Bibby and Lloyd 1977, Ford et al. 1987, Ford 2006), and distance the carcasses would need to drift to reach the shoreline (Martin et al. 2020).

The proportion of mortalities that were lost at sea during this outbreak is unknown but is likely to be significant and to have varied regionally in eastern Canada. As previously mentioned, the dominant currents in eastern and northeastern Newfoundland (Wu and Tang 2011) likely act in concert to advect many carcasses floating in that region offshore. In future, adaptation of operational drift modelling tools developed for oil spill response (e.g., (e.g., Paquin et al. 2020, Sutherland et al. 2022) could be used to estimate the grounding probability of simulated particles configured to drift like seabirds. Such information could be used to estimate at-sea losses and thus improve mortality estimates for specific species.

Across eastern Canada, seabird and sea duck colonies were completely (66), partially (29), and/or incidentally (46) surveyed by government and academic biologists. All information about observed mortalities on colonies has been collated, however, there are important considerations when interpreting our colony-related data. Specifically, the detectability of any mortality at a colony depends on the magnitude of the outbreak, the timing, method and purpose of the survey, and species-specific traits including nesting habitat and whether birds are likely to return to the colony when sick. It is likely that mortality at the colony was easier to detect among surface nesting species (e.g., gulls, terns, pelicans, cormorants, and gannets) than among burrow nesting birds (e.g., puffins and storm-petrels) or among cliff nesting birds (e.g., murres and kittiwakes) which may have difficulty remaining on ledges when sick or dead. Even with surface nesting species like gannets, we experienced challenges in distinguishing between fresh dead birds and decomposing dead birds when enumerating mortalities from aerial photographs when colonies were visited multiple times. Ultimately, the absence of detections of sick or dead birds at the colonies should not be taken as evidence of absence of mortality and population surveys in subsequent years will provide the best measure of any population-level impacts.

### Reflections and recommendations

During the HPAI outbreak, representatives from ECCC, the Canadian Wildlife Health Cooperative, provincial/territorial government agencies, other federal departments (Canadian Food Inspection Agency, Public Health Agency of Canada, Parks Canada, and Indigenous Services Canada), and Indigenous and academic partners worked together to respond to the HPAI outbreak in a One Health Approach (Giacinti et al. 2023). Similarly, communication and collaboration were the cornerstone of the response to the mass mortality event in eastern Canada. Significant efforts were made by all parties to support mortality data collation efforts, however data quality varied across jurisdictions. The magnitude, duration, and geographic scope of the HPAI outbreak and resulting mass mortality event in eastern Canada was unexpected and unprecedented for this region. Similar large-scale events have occurred across the globe in recent years, with seabirds being among the highly vulnerable to HPAI-related mortality. In all cases, estimating total mortality is a logistical challenge.

Beached bird surveys have become an international practice for documenting and monitoring the impacts of various sources of mortality (Camphuysen and Heubeck 2001, Wilhelm et al. 2009, Jones et al. 2023). Given the apparent vulnerability of seabirds and sea ducks to HPAI 2.3.4.4b, standardized beached bird surveys could provide valuable information if implemented in areas of anticipated disease outbreaks or in areas when outbreaks are detected early. They could improve the enumeration of large mortality events by providing information on the onset, duration, and magnitude of HPAI mortality. Beached bird surveys could also provide valuable information on species composition and age classes as well as improve access to fresh carcasses for testing, necropsy, and early confirmation of HPAI. Where the resources to establish beached bird surveys or collating mortality reports is limited, band recoveries (Johnston et al., in prep) and or citizen science data from iNaturalist (e.g., Bartolotta et al. 2023) may provide an inexpensive approach to collect information on mortality events.

## Conclusion

Within months of the first positive detection of HPAI H5N1 2.3.4.4b virus in North America, this virus sparked an outbreak that caused a mass mortality event of unprecedented magnitude and duration and at least forty thousand wild birds across 45 species were reported as sick or dead. Most of the mortalities were among Northern Gannets, Common Murres and Common Eiders, although unusual levels of mortality were also reported in gulls, cormorants, Atlantic Puffins, Black-legged Kittiwakes, Razorbills, and terns. Based on our assessment, it is probable that Northern Gannets and Common Eiders will experience population-level impacts in eastern Canada, whereas such impacts are not anticipated for Common Murres, due to their large breeding populations. We recommend that the breeding populations of these three species in eastern Canada be monitored closely in the years that follow to assess the long-term impacts of HPAI mortality and support conservation decisions. Beached bird surveys could improve the enumeration of large mortality events if implemented in areas where repeated outbreaks are expected and/or ongoing.

## Acknowledgements

Most reported mortalities can be traced back to engaged members of the public, and we offer our sincere thanks to all individuals who shared their observations. Numerous individuals and organizations have been instrumental in completing this research, and their collective contributions have played a pivotal role in shaping the research outcomes (Appendix S4). Special thanks to Campbell Bowser, who’s assistance improved this study. Funding for this research was provided by Environment and Climate Change Canada, NSERC, and Memorial University of Newfoundland.

## Author Contributions

**S. Avery-Gomm:** Conceptualization, Data curation, Formal analysis, Funding acquisition, Investigation, Methodology, Project administration, Software, Supervision, Validation, Visualization, Writing - original draft, Writing - review & editing. **T. Barychka:** Conceptualization, Data curation, Formal analysis, Investigation, Methodology, Software, Validation, Visualization, Writing - original draft, Writing - review & editing. **M. English:** Conceptualization, Data curation, Investigation, Methodology, Writing - review & editing. **R. Ronconi:** Conceptualization, Investigation, Methodology, Writing - review & editing. **S. I. Wilhelm:** Conceptualization, Investigation, Methodology, Writing - review & editing. **J.-F. Rail:** Conceptualization, Data curation, Investigation, Methodology, Writing - review & editing. **T. Cormier:** Data curation, Investigation, Methodology, Writing - review & editing. **M. Beaumont:** Conceptualization, Investigation, Methodology. **C. Bowser:** Software, Validation, Visualization, Writing - review & editing. **T. V. Burt:** Conceptualization, Data curation, Investigation, Methodology, Writing - review & editing. **S. Collins:** Conceptualization, Data curation, Investigation, Methodology, Writing - review & editing. **S. Duffy:** Investigation, Methodology**. J. A. Giacinti:** Data curation, Investigation, Methodology, Writing - review & editing. **S. Gilliland:** Investigation, Methodology, Writing - review & editing. **J.-F. Giroux:** Data curation, Investigation, Methodology, Writing - review & editing. **C. Gjerdrum:** Conceptualization, Investigation, Methodology, Writing - review & editing. **M. Guillemette:** Investigation, Methodology. **K. E. Hargan:** Investigation, Methodology. **M. Jones:** Conceptualization, Data curation, Investigation, Methodology. **A. Kennedy:** Conceptualization, Investigation, Methodology. **L. Kusalik:** Data curation, Software, Validation. **S. Lair:** Conceptualization, Data curation, Investigation, Methodology, Writing - review & editing. **A. S. Lang:** Investigation, Methodology, Writing - review & editing. **R. Lavoie:** Investigation, Methodology. **C. Lepage:** Data curation, Investigation, Methodology, Writing - review & editing. **G. McPhail:** Conceptualization, Data curation, Investigation, Methodology, Writing - review & editing. **W. A. Montevecchi:** Conceptualization, Data curation, Investigation, Methodology, Writing - review & editing. **G. J. Parsons:** Investigation, Methodology, Writing - review & editing. **J. F. Provencher:** Investigation, Methodology, Writing - review & editing. **I. Rahman:** Investigation, Methodology, Writing - review & editing. **G. J. Robertson:** Conceptualization, Investigation, Methodology, Writing - review & editing. **Y. Seyer:** Data curation, Investigation, Methodology, Writing - review & editing. **C. Soos:** Investigation, Methodology. **C. R. E. Ward:** Investigation, Methodology, Writing - review & editing. **R. Wells:** Conceptualization, Investigation, Methodology. **J. Wight:** Investigation, Methodology, Writing - review & editing.

## Notes

https://doi.org/10.6084/m9.figshare.24856869

### Competing Interest Statement

The authors have declared no competing interest.

