## Appendix S4 for "Wild bird mass mortalities in eastern Canada associated with the Highly Pathogenic Avian Influenza A(H5N1) virus, 2022"

### Appendix S2: Supporting details for double count analysis.

**Table S1:** Species groups and species treated as interchangeable during the double count analysis since not all observers will be able to distinguish between similar-looking species.

| **Group Name** | **Species (Common Names)** |
|---|---|
| Gulls | Unknown Gull, Herring Gull, Great Black-backed Gull, Iceland Gull, Glaucous Gull, Ring-billed Gull, Black-headed gulls Laughing Gulls, Lesser Black-backed Gull, Bonapart Gull, Unknown Gull - Large white-headed gulls (Larus *sp.*) |
| Terns | Arctic Tern, Common Tern, Roseate Terns, Unknown Tern |
| Cormorants | Double-crested Cormorant, Great Cormorant, Unknown Cormorant |

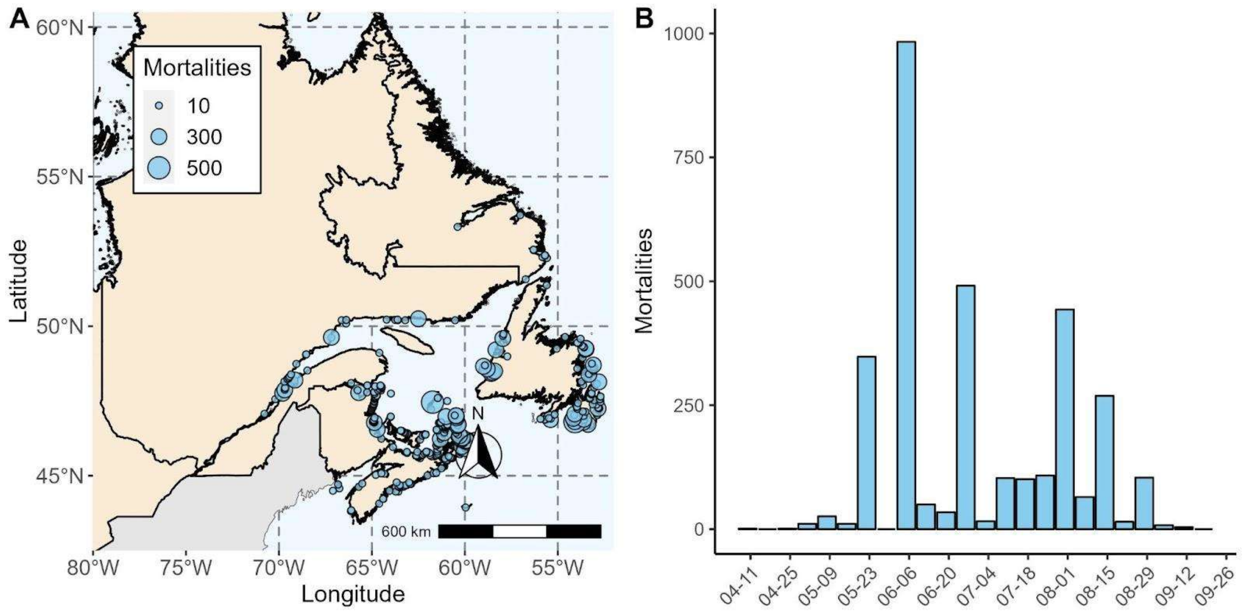


**Figure S1.** The (A) spatial and (B) temporal distribution of mortalities and morbidities identified as double counts in Scenario B (±1 km, ±1 day), and are thus excluded from subsequent summaries of HPAI-linked mortality. Most double counts that were identified by the analysis occurred in Newfoundland and Labrador and Nova Scotia.
