## Appendix S1 for "Wild bird mass mortalities in eastern Canada associated with the Highly Pathogenic Avian Influenza A(H5N1) virus, 2022"

### Appendix S1: Methods for iNaturalist and eBird mortality data

PREPRINT

**Section S1:** iNaturalist - Downloading data and extracting mortality reports.

Citizen sciewatermarnce platforms such as iNaturalist, host a large collection of observations of both flora and fauna contributed by a community of users that can be used to supplement large datasets like the one we used for this report. Extracting data from iNaturalist to target specific questions is typically achieved through the creation of 'projects', a feature within the platform. Projects filter through all observations based on a set of required criteria outlined by the creator of the project. Depending on the intent of the project, the desired observations are presented in a clean, user-friendly interface within iNaturalist or can be exported from the platform for further analysis. To export data from iNaturalist, visit https://www.inaturalist.org/observations/export.

We looked to iNaturalist to find additional reports of dead birds that may have been related to HPAI, within a specified time-period and region. However, recording observations of dead birds (or any wildlife) requires users to re-visit their original observation using a web browser version of the platform and manually edit the submission to reflect the individual as 'dead'. We discovered that some users of iNaturalist were reporting high volumes of dead birds but were unaware they needed to edit each observation. Therefore, we created two projects with different sets of criteria to ensure we included as many dead bird reports as possible.

*Part 1 of 2: Dead Birds in Eastern Canada iNaturalist Project:*

The first project, "**HPAI | Dead birds in Eastern Canada during the 2022 HPAI outbreak**" identified birds that were tagged as dead in Quebec, New Brunswick, Nova Scotia, PEI, Newfoundland and Labrador, and Saint Pierre et Miquelon between April 1 and September 30, 2022, and included a photo of the bird (Table A1). We exported this dataset, and this resulted in 457 records of dead birds.

This iNaturalist project is accessible here: [https://www.inaturalist.org/projects/hpai-dead-birds-in-eastern-canada-during-the-2022-hpai-outbreak](https://www.inaturalist.org/projects/hpai-dead-birds-in-eastern-canada-during-the-2022-hpai-outbreak). Note that the online data is not an archive and exports may vary if records during this time period were edited on the iNaturalist website by observers.

To refine the dataset, we identified a list of words/phrases that can be used to filter out records that are unlikely to be related to HPAI (i.e., broken bird egg, building strike, car, cat, foot and leg of bird, found head separated from body, foxes, ground nest of 1-2 eggs, hawk, lobster trap piece, missing head, not legally hunted animals, predated, remains of foot, roadkill, stray cat, window, window kill, window strike, hit bus shelter glass). Next, where records were classified as 'Needs ID', photos were downloaded and examined to verify species identification to the lowest possible taxonomic level, and this information was added to the dataset. Here, we also excluded records with photos of bleached bones or singular feathers that were deemed too old to be included in our period of study. If the observation took place on land or water, that was noted and added to the dataset as well. Finally, these records were merged with the master list of wild bird mortalities (i.e., HPAI Mortality Tracker).

To export data from "**HPAI | Dead birds in Eastern Canada during the 2022 HPAI outbreak**", we used the following query field in iNaturalist, and selected 'All' in the tables labelled 'Geo' and 'Taxon Extras'.

*quality_grade=any&identifications=any&projects%5B%5D=dead-birds-in-eastern-canada-during-the-2022-hpai-outbreak&d1=2022-04-1&d2=2022-9-30+Columns+id%2C+observed_on_string%2C+observed_on%2C+time_observed_at%2C+time_zone%2C+user_id%2C+user_login%2C+user_name%2C+created_at%2C+updated_at%2C+quality_grade%2C+license%2C+url%2C+image_url%2C+sound_url%2C+tag_list%2C+description%2C+num_identification_agreements%2C+num_identification_disagreements%2C+captive_cultivated%2C+oauth_application_id%2C+place_guess%2C+latitude%2C+longitude%2C+positional_accuracy%2C+private_place_guess%2C+private_latitude%2C+private_longitude%2C+public_positional_accuracy%2C+geoprivacy%2C+taxon_geoprivacy%2C+coordinates_obscured%2C+positioning_method%*

PREPRINT

*2C+positioning_device%2C+place_town_name%2C+place_county_name%2C+place_state_name%2C+place_country_name%2C+place_admin1_name%2C+place_admin2_name%2C+species_guess%2C+scientific_name%2C+common_name%2C+iconic_taxon_name%2C+taxon_id%2C+taxon_kingdom_name%2C+taxon_phylum_name%2C+taxon_subphylum_name%2C+taxon_superclass_name%2C+taxon_class_name%2C+taxon_subclass_name%2C+taxon_superorder_name%2C+taxon_order_name%2C+taxon_suborder_name%2C+taxon_superfamily_name%2C+taxon_family_name%2C+taxon_subfamily_name%2C+taxon_supertribe_name%2C+taxon_tribe_name%2C+taxon_subtribe_name%2C+taxon_genus_name%2C+taxon_genushybrid_name%2C+taxon_species_name%2C+taxon_hybrid_name%2C+taxon_subspecies_name%2C+taxon_variety_name%2C+taxon_form_name*

*Part 2 of 2. NatureNB birds in Eastern Canada iNaturalist Project:*

The second project was created for a specific group of iNaturalist observers from NatureNB who were conducting beached bird surveys but did not record their observations as 'Dead' in 2022 and therefore these records were not picked up by the project. This project is called "**NatureNB Birds in Eastern Canada**" and holds nearly all the same criteria as the "**HPAI | Dead birds in Eastern Canada during the 2022 HPAI outbreak**" project but was targeted for known, high-volume users, and did not require the observation to be annotated as 'dead' (Table S1).

This iNaturalist project is accessible here: https://www.inaturalist.org/projects/naturenb-birds-in-eastern-canada. Note that the online data is not an archive and exports may vary if records during this time period were edited on the iNaturalist website by observers.

For these observers only, we exported all observations within the study window (N = 826), and we downloaded all associated images to examine and identify observations where birds were visibly dead. While sorting through the observations for dead birds only, species were identified to the lowest possible taxonomic level in cases where iNaturalist observations were not classified as "Research Quality" and required ID confirmation. Next, we identified records where the

PREPRINT

description field contained words used to exclude observations unlikely to be HPAI-related (i.e., roadkill, window strikes). After all unnecessary records were excluded, these records were added to the complete mortality dataset.

To export data from "**NatureNB Birds in Eastern Canada**", we used the following query and selected 'All' in the tables labelled 'Geo' and 'Taxon Extras'.

*quality_grade=any&identifications=any&projects[]=naturenb-birds-in-eastern-canada&d1=2022-04-1&d2=2022-9-30 Columns id, observed_on_string, observed_on, time_observed_at, time_zone, user_id, user_login, user_name, created_at, updated_at, quality_grade, license, url, image_url, sound_url, tag_list, description, num_identification_agreements, num_identification_disagreements, captive_cultivated, oauth_application_id, place_guess, latitude, longitude, positional_accuracy, private_place_guess, private_latitude, private_longitude, public_positional_accuracy, geoprivacy, taxon_geoprivacy, coordinates_obscured, positioning_method, positioning_device, place_town_name, place_county_name, place_state_name, place_country_name, place_admin1_name, place_admin2_name, species_guess, scientific_name, common_name, iconic_taxon_name, taxon_id, taxon_kingdom_name, taxon_phylum_name, taxon_subphylum_name, taxon_superclass_name, taxon_class_name, taxon_subclass_name, taxon_superorder_name, taxon_order_name, taxon_suborder_name, taxon_superfamily_name, taxon_family_name, taxon_subfamily_name, taxon_supertribe_name, taxon_tribe_name, taxon_subtribe_name, taxon_genus_name, taxon_genushybrid_name, taxon_species_name, taxon_hybrid_name, taxon_subspecies_name, taxon_variety_name, taxon_form_name Columns id, observed_on_string, observed_on, time_observed_at, time_zone, user_id, user_login, user_name, created_at, updated_at, quality_grade, license, url, image_url, sound_url, tag_list, description, num_identification_agreements, num_identification_disagreements, captive_cultivated, oauth_application_id, place_guess, latitude, longitude, positional_accuracy, private_place_guess, private_latitude, private_longitude, public_positional_accuracy, geoprivacy,*

PREPRINT

*taxon_geoprivacy, coordinates_obscured, positioning_method, positioning_device, species_guess, scientific_name, common_name, iconic_taxon_name, taxon_id*

Keywords in the Comment field that were used to exclude iNaturalist reports of dead birds from inclusion in the HPAI mortality dataset include: Broken bird egg, Building strike, Car, Cat, Foot and leg of bird, Found head separated from body, foxes, Ground nest of 1-2 eggs, hawk, Lobster trap piece, Missing head, not legally hunted animals, Predated, Remains of foot, Roadkill, Stray cat, Window, Window kill, Window strike, Hit bus shelter glass

**Table S1.** Settings for Projects used to filter mortality data from iNaturalist

| **Project Name** | Dead birds in Eastern Canada during the 2022 HPAI outbreak | NatureNB Birds in Eastern Canada |
|---|---|---|
| **Taxa** | Aves | |
| **Location** | New Brunswick<br>Newfoundland and Labrador<br>Nova Scotia<br>Prince Edward Island<br>Québec<br>Saint Pierre et Miquelon | |
| **Users** | Any | ahebert<br>audree_b<br>lewnanny_richardson |
| **Projects** | Any | |
| **Quality Grade** | Research Grade, Needs ID | |
| **Media Type** | Photo | |
| **Date** | April 1, 2022 – September 30, 2022 | |
| **Annotation** | Dead | Any |

PREPRINT

**Section S2**: eBird - Downloading data and extracting mortality reports

*Part 1 of 1: Obtaining data and extracting mortality reports*

eBird is an open-source online platform used by citizen scientists to report wild bird observations. Structured surveys such as colony surveys can also be submitted. Information on dead or sick birds is noted by observers while submitting other bird sightings, using the 'Trip Comments' and/or 'Species Comments' fields. Observers are not supposed to submit observations of dead or sick birds; however, we found observations of dead or sick birds were sometimes submitted.

eBird observations from April 1 to September 30, 2022 were downloaded from eBird website (https://ebird.org/home) . Because of the large file sizes, each province's data was downloaded and processed separately, in two batches: between April 1 and 30th June, and between August 1 and September 30. The updated eBird data is available for download after the 15th of each month.

A list of keywords in English and French was compiled, including words such as 'dead, sick, neurologic'. Downloaded datasets were processed in R version 4.2.2. Keyword hits were used to find any observations of potentially dead or sick birds in 'TripComments' and 'SpeciesComments' fields of the downloaded datasets. As Trip and Species Comments could be repeated multiple times within a dataset (for example, when there were multiple bird sightings by the same observer within a trip), only unique observations with distinct comments were retained.

All observations containing keywords were compiled and processed manually to identify records referring to potentially sick or dead birds. Manual processing involved reading through the Trip and Species Comments, removing any erroneous hits (e.g., referring to 'dead tree', 'dead calm', etc.), recording the number of potentially sick or dead birds, the species names, where they were observed (Land, Water, Colony), age and sex (if reported) and whether any birds have been collected and/or sent for analysis.

PREPRINT

Avery-Gomm, S. et al., 2023. Wild bird mass mortalities in eastern Canada associated with the Highly Pathogenic Avian Influenza A(H5N1) virus, 2022. bioRxiv preprint.

Keywords that were used to identify HPAI-related observations in eBird for inclusion in the HPAI mortality dataset include: dead, dying, death, mortality, deceased, sick, diseased, influenza, neurologic, carcasses, morbid, morbidity, viruses, viral, infection, head tilt, paralysis, seizures, diarrhea, weakness, drooping, distress, mort, mourant, décès, mortalité, décédé, malade, maladie, neurologique, neuro, malade, morbide, morbidité, viral, infecté, infecter, qui tombe, torsion, encerclement, traumatisme, éternuement, décoloration, difficile

PREPRINT
