## Appendix S2 for "Wild bird mass mortalities in eastern Canada associated with the Highly Pathogenic Avian Influenza A(H5N1) virus, 2022"

### Appendix S3: Details on reported mortality for additional species: gulls, cormorants, Atlantic Puffins, Black-legged Kittiwakes, Razorbills, and terns.

Although the largest number of reported mortalities were among Northern Gannets, Common Murres, and Common Eiders, gulls, cormorants, Atlantic Puffins, Black-legged Kittiwakes, Razorbills, and terns also suffered unusual mortality during the HPAI H5N1 2.3.4.4b outbreak.

<u>Gulls</u>

In total, 2,326 gulls were reported sick or dead (Figure S1). Most were Herring Gulls (*n* = 1,030), unknown gulls (*n* = 745), and Great Black-backed Gulls (*n* = 522), with smaller numbers of Ring-billed Gulls (*n* = 26) and Iceland Gulls (*n* = 3). Approximately 66% of the reports came from colonies and 34% were reported on land or at sea. In the St. Lawrence Estuary in QC, mortalities were reported on Île aux Pommes and Île Blanche in May and June (10 Great Black-backed Gulls, 14 Herring Gulls, 218 unknown gulls) and on Île Bicquette in June (85 Herring Gulls, 70 Great Black-backed Gulls).

Mortalities were also reported on Kent Island, NB (182 Herring Gulls, July - September) and Bear Island, NS (302 Great Black-backed Gulls and 302 Herring Gulls, early August - early September). Observations from Bear Island indicate the remains were old and mortalities from an HPAI outbreak likely occurred between August 2 and September 3. One dead Great Black-back Gull was reported on Machias Seal Island, where large gull nesting attempts are actively discouraged to protect tern nesting habitat.

Gull mortalities were reported off colonies from early April until late September, peaking in June and early August. Mortalities (both on and off colonies) were reported in all provinces. A total of 784 gull mortalities were reported in QC, 865 were reported in NS, 383 were reported in NB, 288 were reported in NL (including 58 in southern Labrador), and 6 gulls were reported in PE. Peaking in Quebec in May and June, most reports of gull mortalities came from the island of



PREPRINT

Newfoundland through July and August, with birds reported along the coastlines of the Burin, Avalon, and Bonavista Peninsulas.

<ins>Cormorants</ins>

Cormorants were the fifth most reported species group (*n* = 981; Figure S2). This group includes Double-crested Cormorants (*n* = 813) and unknown cormorants (*n* = 168). The majority of the mortalities occurred in colonies (71.2%), with ~600 of those occurring at four colonies in Miramichi Bay, NB, in early June.

Cormorant mortalities were reported between mid-May and late September, with a peak in June. A total of 660 mortalities were reported in NB, 281 were reported in QC, 32 were reported in NS, 6 were reported on the island of Newfoundland (none in Labrador), and 2 cormorants were reported in PE.

<ins>Atlantic Puffins</ins>

Atlantic Puffins were the sixth most reported species, with all 282 mortalities reported on the island of Newfoundland between mid-July and September (Figure S3). Most reports came from eastern Newfoundland, with large numbers reported on the Burin, Avalon, and Bonavista Peninsulas. Very few mortalities were observed at the breeding colonies that were visited (*n* = 6), and this low number of mortalities could not be distinguished from background mortality rates.

<ins>Black-legged Kittiwakes</ins>

Black-legged Kittiwakes were the seventh most reported species (*n* = 251; Figure S4). Mortalities were reported between June and September, with mortality reports peaking in late July. Most mortalities were reported in NL (*n* = 222, including 5 in Labrador), with smaller numbers reported in QC (*n* = 27) and NB (*n* = 2). Most mortalities occurred off of colonies (90%). Most Black-legged Kittiwake mortalities reported in NL (*n* = 203 of 222 birds) likely

PREPRINT

originated from colonies in the Witless Bay and Cape St. Mary's Ecological Reserves, which host ~24,000 and ~8,800 breeding kittiwakes, respectively; S. Wilhelm, pers. comm.). A survey of a Black-legged Kittiwakes breeding at Gull Island in Witless Bay Ecological Reserve revealed two mortalities; however, this species builds their nests on cliff ledges, so they may be more prone to being observed dead in the water around colonies than on the colony itself. A small outbreak (*n* = 20) was also reported in the waters below a kittiwake colony on Île Brion, in Îles-de-la-Madeleine (~8,420 breeding birds in 2017) in the beginning of August. (J.-F. Rail, pers. comm.).

Razorbills

Razorbills were the eighth most reported species (*n* = 119; Figure S5). Most mortality was reported in NL (*n* = 82, including 3 in Labrador), with smaller numbers reported in QC (*n* = 34) and NB (*n* = 3). Most occurred off colonies (90.8%) between mid-May and late August. The colony mortalities (*n* = 11) were reported in QC's Migratory Bird Sanctuaries on the North Shore of the Gulf of St. Lawrence, on Île aux Pommes, Île aux Oeufs, Pot du Phare and Gros Pot in the St. Lawrence Estuary. No mortalities were observed during a boat survey of Gull Island in Witless Bay Ecological Reserve, Newfoundland on July 22, 2022, and during foot surveys of Machias Seal Island and Green Rock, NB in June 2022.

Terns

In total, 74 terns were reported between the end of May and the beginning of September, with a peak in mortalities during June (Figure S6). This group includes unknown terns (*n* = 51), Arctic Terns (*n* = 13) and Common Terns (*n* = 10). Roseate Terns were not reported in the dataset but were present at sites where other species tested positive for the HPAI virus. Most mortalities were reported in NB (*n* = 36) and NS (*n* = 35), with smaller numbers reported in NL (*n* = 3, including 2 in Labrador). Most reported mortality occurred in colonies (86.5%) in Shelburne, NS (*n* = 33) and Tern Islands in Kouchibouguac National Park, northeastern NB (*n* = 29).

PREPRINT

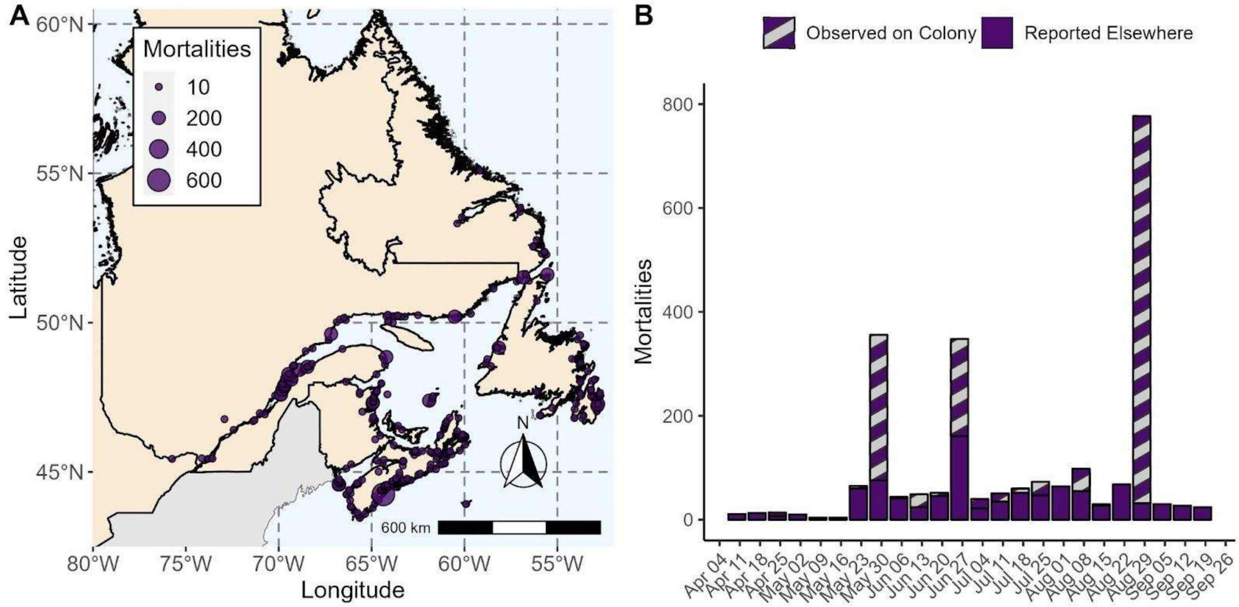


**Figure S1.** The spatial (A) and temporal (B) distributions of HPAI-linked mortality and morbidity for gulls, with double counted birds removed (Scenario B: ±1 km, ±1 day). Herring Gull, Great Black-backed Gull and Unknown Gull are represented in this group, along with a small number of other gulls.

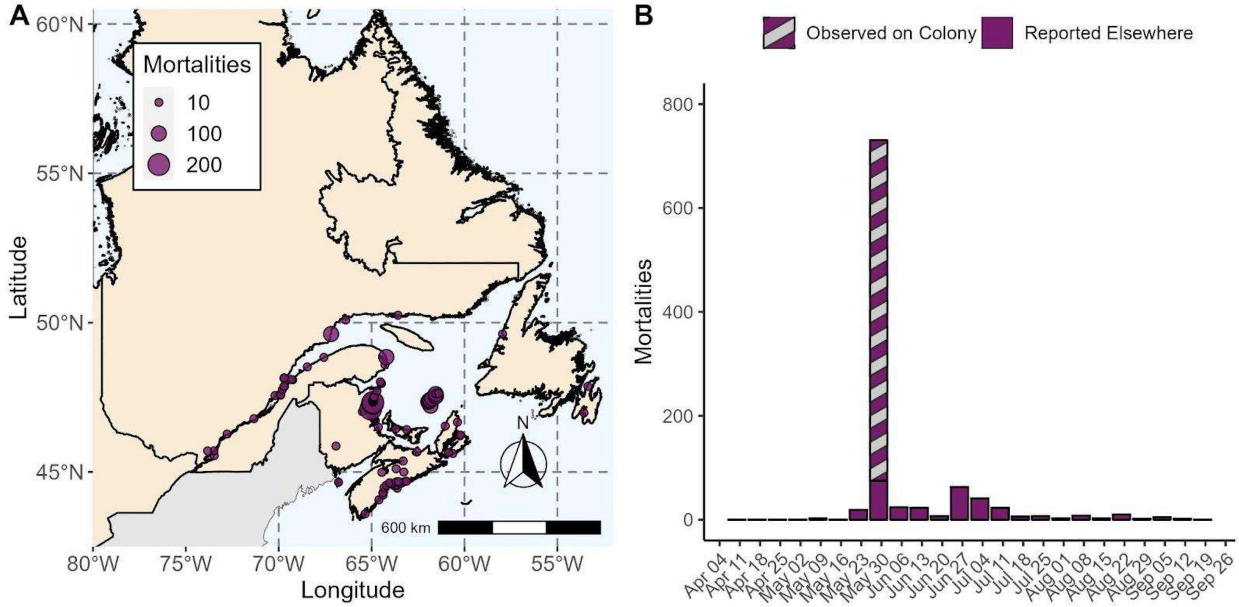

**Figure S2.** The (A) spatial and (B) temporal distribution of HPAI-linked mortality and morbidity for cormorants, with double counted birds removed (Scenario B: ±1 km, ±1 day). Double-crested Cormorant, Great Cormorant and Unknown Cormorant are represented in this group.

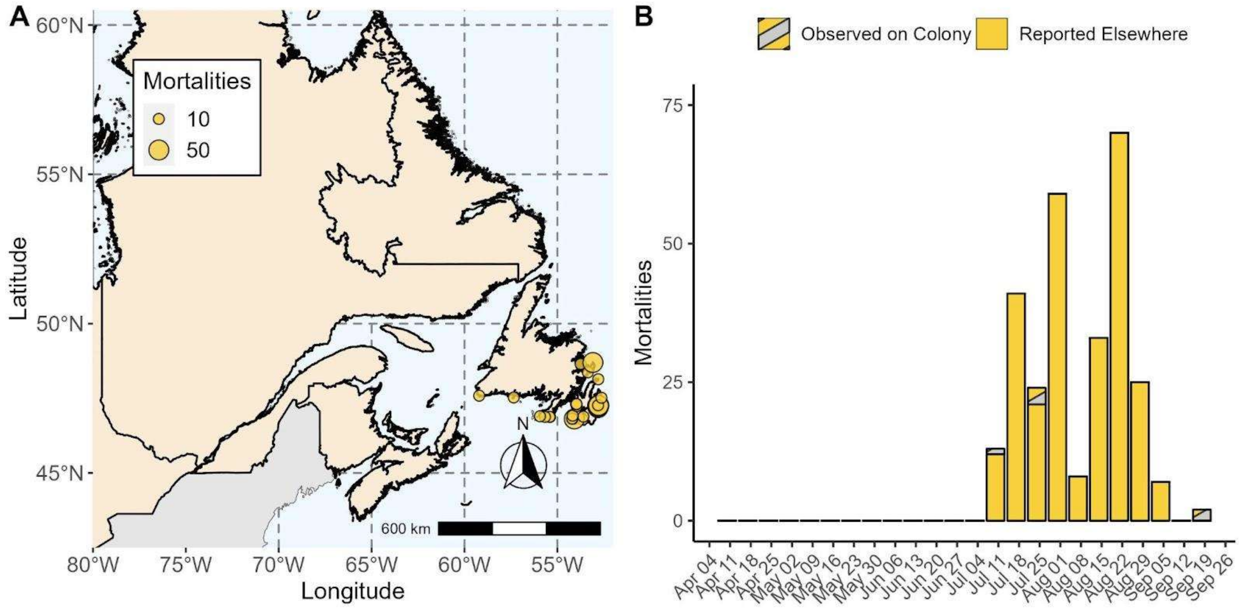


**Figure S3.** The (A) spatial and (B) temporal distribution of HPAI-linked mortality and morbidity for Atlantic Puffins, with double counted birds removed (Scenario B: ±1 km, ±1 day).

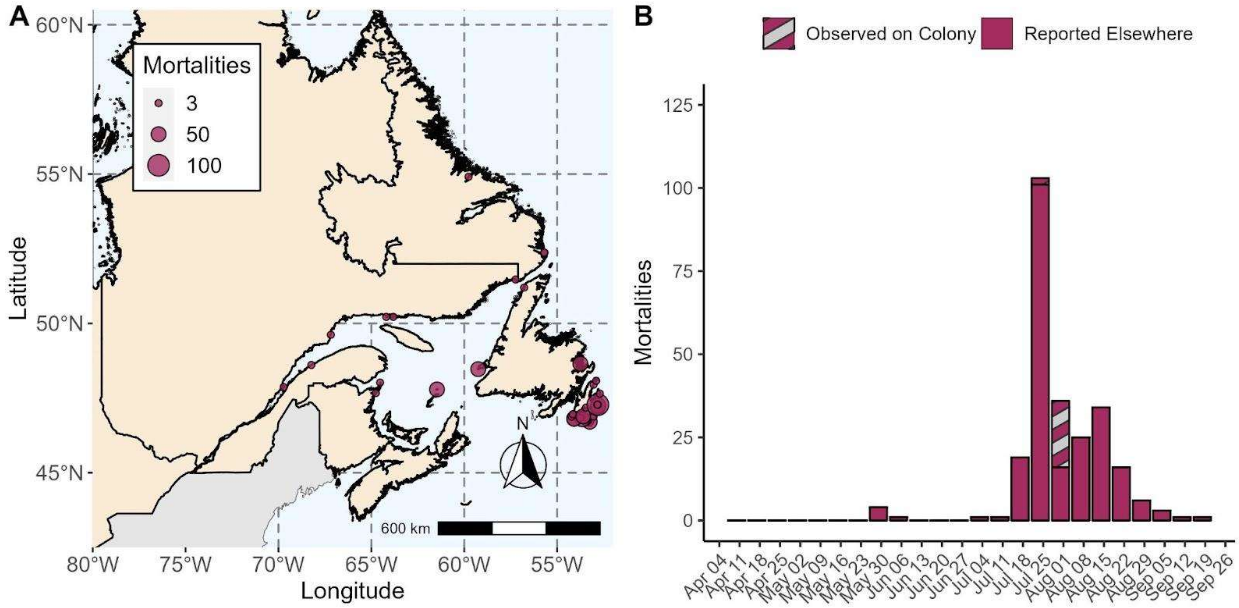


**Figure S4.** The (A) spatial and (B) temporal distribution of HPAI-linked mortality and morbidity for Black-legged Kittiwakes, with double counted birds removed (Scenario B: ±1 km, ±1 day).

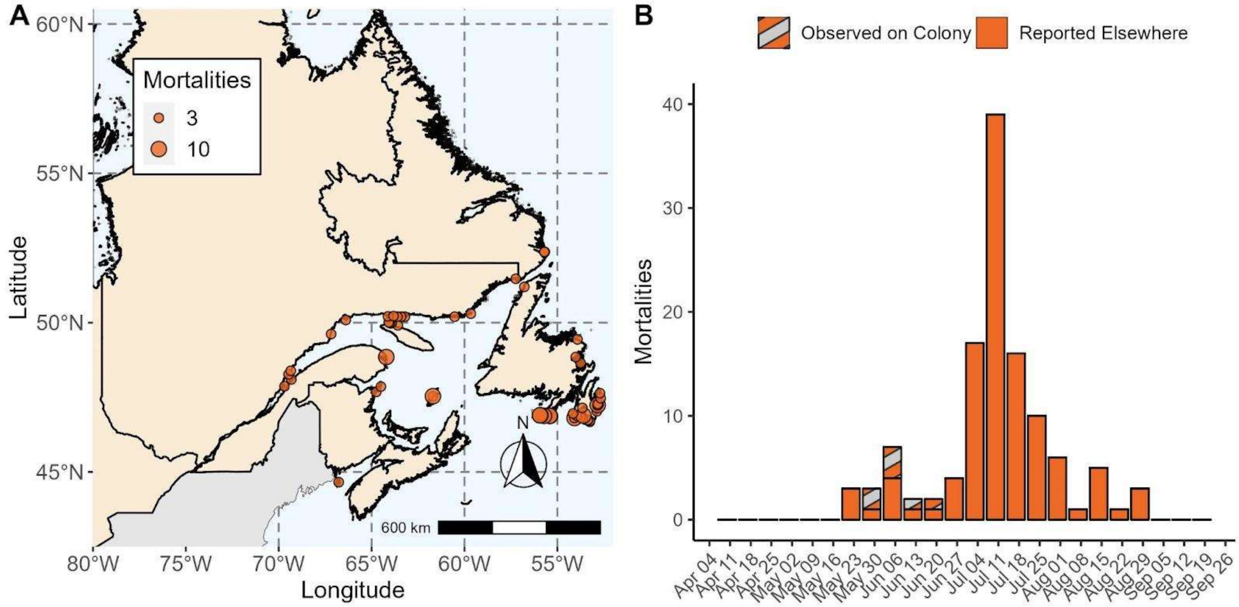


**Figure S5.** The (A) spatial and (B) temporal distribution of HPAI-linked mortality and morbidity for Razorbills, with double counted birds removed (Scenario B: ±1 km, ±1 day).

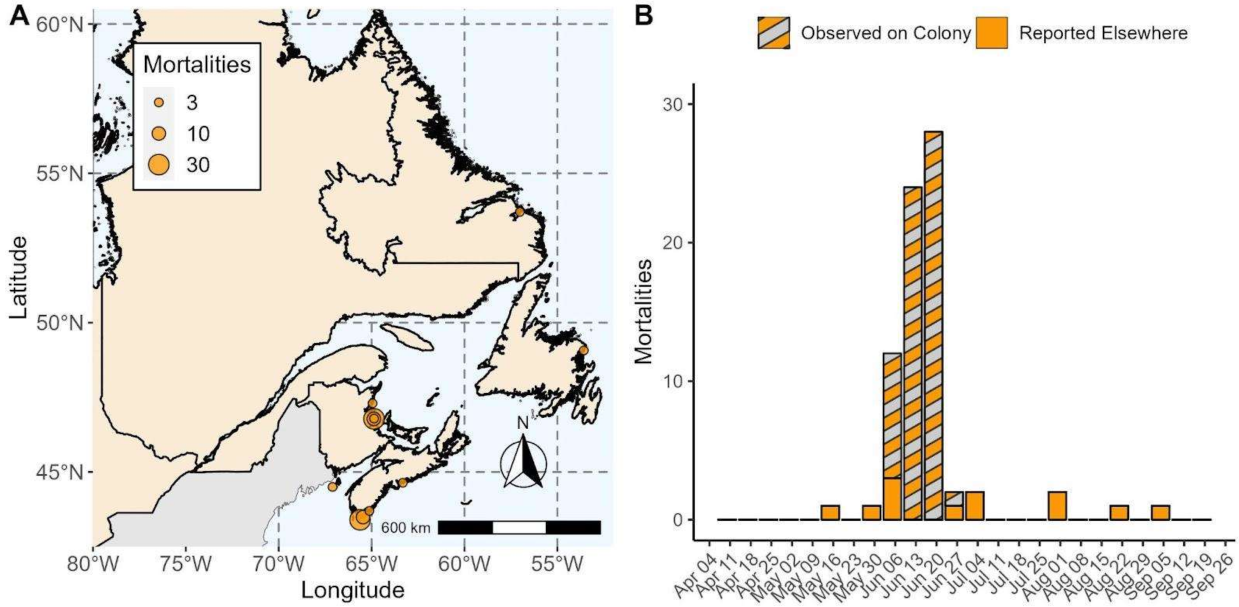


**Figure S6.** The (A) spatial and (B) temporal distribution of HPAI-linked mortality and morbidity for terns, with double counted birds removed (Scenario B: ±1 km, ±1 day). Arctic Tern, Common Tern and Unknown Tern are represented in this group.
