## Appendix S3 for "Wild bird mass mortalities in eastern Canada associated with the Highly Pathogenic Avian Influenza A(H5N1) virus, 2022"

### Appendix S4: Additional Acknowledgements

Most reported mortalities can be traced back to engaged members of the public, and we offer our sincere thanks to all individuals who shared their observations. Numerous individuals from Indigenous, federal, provincial, and municipal governments as well as non-governmental have been instrumental in the completion of this research. Thank you.

**Indigenous partners:**
This research was conducted in what is now known as eastern Canada on the traditional land of the Mi’kmaq First Nation, the Wolastoqiyik (Maliseet) First Nation, the Passamaquoddy people, the Beothuk peoples, the Inuit of Nunatsiavut, the Inuit of NunatuKavut and the Innu of Nitassinan. We would like to acknowledge the support provided by members of the Innu Nation, the Miawpukek First Nation (MFN), the Nunatsiavut Government, the NunatuKavut Community Council, and the Qalipu First Nation.

**Government Agencies and NGOs:**
We would like to acknowledge the invaluable support provided by staff from the following organizations:

Canadian Wildlife Health Cooperative (CWHC - Altantic, CWHC - Quebec), Ministère de la faune et des parcs, Ministère de l’agriculture et de l’alimentation, Newfoundland and Labrador Department of Fisheries, Forestry and Agriculture (including Regional Services, Animal Health Division, Natural Areas Division and Wildlife Division), Nova Scotia Department of Natural Resources and Renewables (Wildlife Division), Prince Edward Island Department of Environment, Water and Climate Change, Forests, Fish and Wildlife Division, Municipality of the Magdalen Islands (Les Iles-de-la-Madeleine), ACAP Cape Breton, Birds Canada, Nature NB, Canadian Parks and Wilderness Society - The Newfoundland and Labrador Chapter (CPAWS - NL), Indian Bay Ecosystems Initiative, Rock Wildlife Rescue, Société protectrice des eiders de l’estuaire, Société Provancher, Parc National de l’Île-Bonaventure-et-du-Rocher-Percé, Environment and Climate Change Canada, Parks Canada (including Mingan National

PREPRINT

Park, Prince Edward Island National Park, Kouchibouguac National Park, Cape Breton Highlands National Park, Saguenay National Park, Kejimkujik National Park, Forillon National Park), Fisheries and Oceans Canada, Transport Canada's National Aerial Surveillance Program

**Individuals:**

Our deepest appreciation goes to the following individuals who contributed mortality data to this study:

April Hedd, Alyssa Hunter, Anabelle Hebertz, Ariane Massé, Audrée Benoit, Aurore Perrot, Beverly Dawe, Blair Adams, Bob Petrie, Brad Potter, Brad Romaniuk, Charlene Kippenhuck, Chris Baldwin, Chris Mooney, Christine Lepage, Christopher Poole, Chuck Porter, Cynthia Pekarik, Daniel Gallant, Daren Sheppard, Darrin Reid, Dave Fifield, Dave McRuer, Denise Maillet, Drew Hutchinson, Donna Hurlburt, Elena Haratsaris, Eliza-Jane Morin, Erin Muntz, Erin Ramsay, Erin Shea, François Lapointe, Frederic Dwyer-Samuel, Frédérick Lelièvre, Gabrielle Dimitri-Masson, Gail Davoren, Garry Gregory, Gerry Pasteen, Gregoy Jeddore, Harry Worthman, Jason Dicker, Jayne Boutilier, Jean-Simon Richard, Jeannine Winkle, Jim Rudderham, Jonathan Strickland, Josh Cunningham, Karen Gosse, Katharine Studholme, Kathleen Aikens, Kaylene Stagg, Kevin Craig, Kim Huskins, Leighann Hartnett, Lewnanny Richardson, Linda Redmond, Mathieu Cote, Michelle Saunders, Megan Baker, Megan Fortune, Michael Hannoford, Michael Brown, Paul Kozie, Reg Delorey, René Nault, Shavonne Meyer, Steven Stewart, Suzanne Dooley, Terry Power, Thibaud Durbecq, Tina Leonard, Todd Hollet, Tony Power, Trevor Thompson, Tricia Fleming, Wesley Morgan

PREPRINT
